# Data-driven dynamic modelling identifies polyploidisation as key process in cell cycle progression upon DNA damage

**DOI:** 10.1101/2025.08.14.670293

**Authors:** Elsje J. Burgers, Muriel M. Heldring, Lukas S. Wijaya, Tamara Y. Danilyuk, Jiahang Su, Sebastiaan J. Molenaar, Peter Bouwman, Sylvia E. Le Dévédec, Bob van de Water, Joost B. Beltman

## Abstract

Chemotherapeutic agents often cause DNA damage in order to kill fast-dividing cancer cells or disrupt their proliferation. Therefore, understanding the interplay between DNA damage and cell cycle progression is highly relevant for understanding cancer cell behaviour. An important regulator is transcription factor p53, primarily known for its function to maintain genomic stability, regulate transient and permanent cell cycle arrest and apoptosis. Activated p53 transcriptionally regulates the expression of many proteins, among which are MDM2, p21 and BTG2. MDM2 functions as a direct inhibitor of p53 by targeting it for ubiquitination. The proteins p21 and BTG2 are known for their regulatory function in G1 and G2 cell cycle arrest. Using HepG2-FUCCI cells, we showed that exposure to cisplatin or etoposide caused a temporary G2 arrest. To study the link between protein expression and cell cycle arrest, we developed a mathematical model in which we integrated a previously established model for the protein expression dynamics of p53, MDM2, p21 and BTG2 with a cell cycle model. This allowed us to determine the importance of p21 and BTG2 in their stimulation of G1 and G2 cell cycle arrest. We found that the protein dynamics could predict the G2 cell cycle arrest in exposed cells, but only in combination with endoreplication, i.e., the alternation of S and G phases without mitosis, resulting in polyploid cells. Our model predicted that the majority of cells endoreplicate upon exposure to high concentrations of cisplatin and most concentrations of etoposide, which we validated with additional time-lapse imaging data in which we could track individual cells. In conclusion, polyploidisation is a generic response of HepG2 cells after treatment with DNA-damaging compounds.

## Introduction

Genotoxic agents are widely used as anti-cancer treatments. Their ability to damage the DNA can cause cell death and disrupt cell proliferation, especially in highly proliferative cells like cancer cells [1]. However, the DNA-damaging drugs can also affect healthy cells and thereby cause severe side effects [2]. Understanding how DNA damage affects the cell cycle can increase our mechanistic understanding of the behaviour of cancer cells and is essential for improving cancer treatments. Mathematical models can help to unravel the quantitative interplay between DNA damage and cell cycle progression and thereby contribute to our knowledge on the effect of genotoxic agents on proliferative cells.

Transcription factor p53 is well known for its function as protector of the DNA, because it responds to DNA damage and transcriptionally regulates DNA-damage signal effectors that are involved in DNA repair, cell cycle arrest, and apoptosis. Nevertheless, the processes that determine whether cells commit to cell cycle arrest or apoptosis are not fully understood. Several studies showed that the activity of p53 as a transcription factor and thereby differential regulation of the expression of genes involved in cell fate processes is dependent on post-translational modifications (reviewed in [3, 4]). In addition, the oligomeric state of p53, i.e., monomeric, dimeric or tetrameric p53 structures, affects the expression of target genes and consequent cell fate decisions [5]. Besides the form of p53 itself, the presence of cofactors is essential for the activation of several genes and thereby the ensuing cell fate. For example, the cell cycle inhibitor p21 is only activated by p53 in the presence of p300 [6] and ASPP protein specifically enhances p53-dependent apoptosis, but not cell cycle arrest, by physically interacting with p53 [7]. Yet another factor that influences cell fate decisions is prior p53-mediated mitochondrial apoptotic priming, which promotes apoptosis in case of p53 reactivation [8]. Finally, the role of the temporal dynamics of p53 in cell fate determination has been extensively studied. Pulsatile p53 dynamics represent a protective mechanism that increases the likelihood of recovery [9, 10], whereas sustained, rapid p53 accumulation leads to cell death [11]. Thus, many factors influence the functionality of p53, the expression of its downstream targets and resulting cell fate.

To understand the p53-dependent regulation of cell behaviour, it will be important to quantify how downstream target gene expression relates to cell fate. The cyclin-dependent kinase (CDK) inhibitor p21 is an important p53 downstream target and is mostly known for its role in cell cycle arrest. During the replicative cell cycle, proteins of the cyclin family drive the progression of one cell cycle phase to the next. In mammalian cells, this is regulated through the expression of the cyclins C and D during G1 phase, cyclin E during G1/S transition, cyclin A during S and G2 phase and cyclin B during mitosis [12], and through subsequent binding and activation of these cyclins to their specific CDK binding partners. The complexes cyclin D-CDK4/6 and cyclin E-CDK1/2 are inhibited by p21, which causes G1 arrest [12, 13]. However, G1 arrest is not solely dependent on p21. The p53-inducible protein BTG2 is also known for its antiproliferative function [14] and can promote G1 cell cycle arrest via downregulation of cyclin D [15]. Moreover, p21 inhibits cyclin A-CDK1/2 complexes and to a lesser extent cyclin B/CDK1 [16], whereas BTG2 is a more potent inhibitor of cyclin B and can cause G2/M arrest and cell death [17]. These findings underline the importance of p21 and BTG2 as regulators of cell cycle progression, yet they do not reveal the relative importance of these effects on the cell cycle.

DNA damage can have other consequences on cell fate besides cell cycle arrest or cell death. Specifically, various forms of DNA damage, such as ionizing radiation treatment and several DNA-damaging chemicals, can also cause endoreplication (also known as endoreduplication or endocycling) [18]. Endoreplication is an alternative cell cycle program which consists only of S and G phases and fully lacks the mitotic phase, resulting in polyploid cells with a single enlarged nucleus [19]. Topoisomerase II (topo II) is thought to be a key player in drug-induced endoreplication, and is required to untangle double-stranded DNA which it cuts and rejoins. [20, 21]. Indeed, Cortés et al. [18] propose that inhibition of topo II hinders the segregation of daughter chromatids and thus prevents mitosis. Apart from topo II inhibitors, other genotoxic agents can also cause endoreplication because DNA modifications might prevent topo II from binding to the DNA, and consequently, DNA separation would also fail [18]. For instance, endoreplication has been observed following cisplatin exposure in PROb cells (rat colon carcinoma) [22], or following alpha radiation in Chinese hamster cells [23].

In this study, we investigated the influence of DNA damage on cell cycle progression upon exposure to cisplatin (CDDP) and etoposide (ETO). To quantitatively describe the relation between p21/BTG2 expression and the cell cycle, we used a previously established ordinary differential equation (ODE) model for the DNA damage signalling pathway and coupled this to a cell cycle progression model that simulates the evolution of the number of cells in different cell cycle phases. We used HepG2 cells with a fluorescent ubiquitination-based cell cycle indicator (FUCCI) construct in combination with live-cell confocal microscopy to measure the temporal dynamics of the number of cells in different cell cycle phases. These data showed that cisplatin-induced DNA damage causes a temporary cell cycle arrest in G2 with subsequent cell cycle continuation and arrest in G1. By calibrating various mathematical models to these data, we found that a transition from G2 to G1 without mitosis was required to explain the cell cycle dynamics following cisplatin exposure. Moreover, our models predicted more frequent occurrence of endoreplicating cells for high concentrations of cisplatin than for low concentrations. We recapitulated similar findings for exposure to etoposide, where the temporary G2 arrest was most profound for low concentrations of the drug. Moreover, our models predicted frequent endoreplication already for low concentrations of etoposide. By tracking the fate of individual cells in additional FUCCI time-lapse imaging we validated the model predictions regarding the important role of polyploidisation for DNA damaging agents, including cisplatin.

## Methods

### Cell culture

Human hepatoma (HepG2) cells were purchased from ATCC - Germany (clone HB8065) and maintained in DMEM high glucose (Fisher Scientific - Bleiswijk, The Netherlands) supplemented with 10% (v/v) FBS (Fisher Scientific - Bleiswijk, The Netherlands), 250 U/ml penicillin and 25 µg/ml streptomycin (Fisher Scientific - Bleiswijk, The Netherlands) in humidified atmosphere at 37 degrees Celsius and 5% CO2/air mixture. For the exposure experiments, we used the cells at a passage number lower than 20 and seeded in Greiner black µ-clear 96-well (Greiner, 655096) or 384-well plates (Greiner, 781866), at 32,000 or 4378 cells per well, respectively.

### Stable FUCCI reporter cell line generation

We generated the HepG2-FUCCI reporter cell line using a lentiviral vector. We generated the lentivirus containing FUCCI plasmid (Addgene, #86849) in HEK293T cells cultured in the same medium as mentioned above. HEK293T cells were seeded in 6 cm dishes at a density of 400,000 cells per dish and reached 60-80% confluency after 24 h. For transfection 24 h after seeding, we prepared a mixture of the viral particle DNA composed of VSV (0.7 µg), GAG (1.3 µg), REV (1.0 µg) and the FUCCI-expressing plasmid (2 µg) in 150 µl serum and antibiotic free medium. Next, we prepared a mixture of polyethylenimine (PEI) (10 µl) in serum and antibiotic free medium (140 µl) and incubated it for 5 minutes at room temperature (RT). Hereafter, we slowly added the DNA mixture, followed by another 15 minutes of incubation time. We added the mixture drop by drop to the HEK293T cells cultured in antibiotic free medium covering the culture surface area. After 24 h, we exchanged the medium with antibiotic free medium. We harvested the virus in the medium by collecting and passing the medium through a 0.45 µm sterile filter to eliminate the cell debris 48 h after the transfection. We performed transduction of the HepG2 cells that had grown in a 6-well plate to 70% confluency. On the day of transduction, we replaced the medium of the cells with 1 ml of the medium containing lentivirus carrying the FUCCI construct and added 1 µl of polybrene to enhance the transduction efficiency. We replaced this medium with culture medium after 24 h of transduction. The HepG2-FUCCI cells were then allowed to proliferate to obtain sufficient cells for further experiments.

### Exposure and imaging

We exposed HepG2-FUCCI cells to chemicals two days after seeding on a 384-wells plate. We replaced the medium with medium containing Hoechst 33342 (1:10000 dilution, 1 mg/mL stock) for nuclear staining 2 h before exposure. After 2 h of incubation with Hoechst, we replaced the medium with medium containing cisplatin (Ebewe, The Netherlands) or etoposide (Sigma Aldrich, E1383). We exposed to 1, 2.5 or 5 µM cisplatin or 0.5, 1, 2.5, 5 or 10 µM etoposide. To take into account biological variability, yet simultaneously minimise technical variability, cells originating from three different flasks were seeded and exposed in triplicate on the same day and on the same plate, generating three biological replicates. Directly after compound exposure, we transferred the plate to a Nikon TiE2000 confocal laser microscope (laser: 540 nm for Cdt1 detection, 488 nm for Geminin detection, and 408 nm for Hoechst nuclear staining detection) equipped with an automated stage and perfect focus system, and started imaging with a Nikon CFI Super Fluor 20x magnification objective (numerical aperture 0.75) at two distinct positions per well (i.e. two technical replicates) for 72 h with one imaging round per 25 minutes.

For time-lapse imaging at high magnification, we followed the same procedures as described above, but generated now four biological replicates and supplemented the medium with Annexin V (AnV) for cell death measurements. Immediately after compound exposure, the plate was imaged using a Nikon ECLIPSE Ti2-E confocal microscope (galvano modality) equipped with lasers at 540 nm (Cdt1), 488 nm (Geminin), 408 nm (Hoechst) and 630 nm (Annexin V), automated stage, and perfect focus system. We imaged with 40x magnification objective (2x optical zoom, numerical aperture 1.15, pinhole size 30) at two distinct positions in each well (i.e. two technical replicates) for 66 hours with one imaging round per 1.67 hours. During imaging, all plates were maintained in a humidified atmosphere at 37 degrees Celsius and 5% CO2 /air mixture.

### Image and data analysis

#### Analysis of FUCCI data

For the 20x magnified FUCCI dataset, we identified the nuclei in each image based on the Hoechst signal with the watershed masked clustering algorithm [24] in ImageJ. The segmentation images together with the confocal microscopy images for Hoechst, Geminin-GFP and Cdt1-RFP were loaded into CellProfiler version 3.1.9. To obtain the Geminin-GFP and Cdt1-RFP intensity per nucleus, we executed a pipeline that used the IdentifyPrimaryObjects and MeasureObjectIntensity modules and the H5CellProfiler method [25] to convert the exported HDF5 files into a data matrix containing quantifications per single cell.

We performed various subsequent data analysis steps on the fluorescence mean intensities within segmented nuclei using R version 4.2.2. Because a small proportion of the Cdt1-RFP signal leaked into the Geminin-GFP emission spectrum, the Geminin-GFP mean intensities were consistently increased with a proportion *c* of the Cdt1-RFP signal. We approximated *c* by obtaining the nucleus with the highest Cdt1-RFP/Geminin-GFP ratio and taking *c* as the ratio between the Geminin and Cdt1 expression for this nucleus. With the formula *G_corr_* = *G* − *c* · *G*, where *G_corr_* is the corrected Geminin intensity, *G* the Geminin intensity and *c* the Cdt1 intensity, we corrected the Geminin-GFP expression. Based on the low expression of Cdt1 and Geminin at the start of imaging, we set the threshold for the intensity below which cells were considered Cdt1- or Geminin-negative at 5% of the maximum expression. Such colourless cells were defined as early G1 or G0 phase cells. Cells with a Cdt1-RFP intensity higher than the threshold were classified as cells in G1 phase, whereas cells with a Cdt1-RFP intensity lower but Geminin-GFP intensity higher than the threshold were categorised as S-G2-M cells. Per measurement time point, we counted the number of cells in each phase and normalised these numbers by dividing them by the total number of cells per image at the start of imaging. We used the mean of the two technical replicates to obtain a single value for each biological replicate.

#### Analysis of magnified FUCCI data

For the 80x magnified FUCCI dataset, we segmented the cells using a U-Net deep neural network. In order to train the network, we first generated a ground truth data set by manually annotating images generated with QuPath [26]. In total, we annotated 64 images: 42 in the Hoechst channel, 11 in the Geminin-GFP channel, and 11 in the Cdt1-RFP channel. All 2D images had a resolution of 1024×1024 pixels. The segmentation model was based on a deep 2D U-Net architecture adapted from the nnU-Net framework [27]. The network consisted of 9 encoder-decoder stages, each containing two convolutional layers with 33 kernels, instance normalisation, and LeakyReLU activations. Feature channels doubled with network depth, starting from 32 and reaching up to 512. Down-sampling was performed using stride-2 convolutions, while up-sampling used transpose convolutions combined with skip connections from the encoder. The input patch size matched the image size (1024×1024), with no spatial rescaling (spacing 1.0x1.0). Z-score normalisation was applied to the input images. The network was trained using dice loss, operating on full-resolution microscopy images. The model performance was evaluated in a five-fold cross-validation fashion. At the end, the model performance achieved 0.90 ± 0.024. The intraobserver variation of the annotation was 0.92, indicating that model performance was saturated.

To analyse nuclear size over time, we only included technical replicates (i.e. well positions) with at least 20 cells at the initial time point, and conditions for which we had data from at least 3 biological replicates. To analyse the change in nuclear size over time, we used the segmentation of the Hoechst channel as described above. The size of each nucleus was determined as the number of pixels comprising the surface area. We combined the nuclei from all technical and biological replicates per condition to analyse the distribution of sizes at several time points.

To quantify the fraction of cells dividing or continuing to G1 without division in the 80x dataset, we manually tracked the individual cells in the same 80x images as used for the population analysis. Each cell that was judged as green (Geminin-GFP positive) by the observer was followed over time and classified once it divided, or became red (Cdt1-RFP positive) or colourless without division. We indicated for the non-dividing cells whether they became red or colourless, but considered them in both cases as cells with skipped mitosis. When determination of the cell fate was impossible, or something else happened (e.g. cell death), the event was classified as “unclear” and excluded from further analysis. Green cells that left the imaging frame without visible colour change or that were still green at the end of the imaging period were also excluded from further analysis. The data of technical replicates was combined, after which we determined the fraction of cells dividing and skipping mitosis for each biological replicate. The average of the fractions for each condition was also calculated.

#### Analysis of DNA damage response imaging data

The image-based protein expression data (of *p*53, *MDM* 2, *BTG*2, *p*21) after cisplatin and etoposide exposure were previously published in [28]. Wijaya et al. [28] exposed a panel of eleven reporter cell lines to eight concentrations of thirty different compounds, and imaged them for more than 60 hours. Next, the images were segmented and the intensity of each reporter was quantified on a single cell level. We processed the etoposide data from this study for the relevant reporters in a similar manner as we previously analysed the data for cisplatin [29]. For full details of the data processing strategy we refer to [29]. Briefly, we used the protein expression data on single cell level and calculated the geometric mean of the GFP expression over all cells per image (i.e. per technical replicate). Next, we calculated the average GFP value of the two technical replicates and performed background subtraction per plate and per time point by subtracting the average of the DMSO controls from the average in etoposide exposure conditions. Finally, we applied min-max normalisation to obtain scaled GFP-intensity values between 0 and 1, and aligned the measurement time points of the three biological replicates with a B-spline function (df = 6, degree = 3) to obtain a readout every 1.5 h, starting at 1 and ending at 71.5 h.

#### Analysis of cell death data

We used the readouts for propidium iodide-positive (PI+) and Annexin V-positive (AnV+) cells from [28] to calculate the fraction of dead cells following cisplatin or etoposide exposure. To minimise variability, we included the PI and AnV data of all eleven reporter cell lines and averaged across all biological replicates. Specifically, we classified a cell as PI+ or AnV+ if the overlap between its nucleus and the AnV or PI fluorescent signal was more than 10% of the nuclear area. Furthermore, we divided the cell count for each replicate by its value at the initial time point to obtain the normalised cell count. For the remainder of the work in the current study we only used concentrations of cisplatin and etoposide for which both the mean fraction AnV+ and PI+ cells did not exceed 0.1 and there was no obvious decrease in the mean normalised cell count (Supplementary Figure 1). This resulted in using data up to 5 µM cisplatin and up to 10 µM etoposide.

### Modelling

#### p53 signalling model

We used the model in [30] as a base for our p53 signalling model to simulate p21 and BTG2 dynamics. In brief, phosphorylation of p53 (*p*_53_) triggered by drug-induced DNA damage (*DD*) causes transcriptional activation of MDM2 (*M*_2_), p21 (*p*_21_) and BTG2 (*B*_2_), while MDM2 increases p53 degradation. Compared to the original model, we removed the scaling and offset of the variables (similar as in [31]) and we simplified the description of p53 by using only one equation instead of separate equations for p53 and phosphorylated p53. Thus, in this simplified description we considered the nuclear p53, which is measured in the imaging data, to be representative for phosphorylated p53. Initially, we optimised the model parameters for the cisplatin and etoposide DNA damage response data simultaneously. In a model extension, we subsequently included *α* as a scaling factor for modelling the etoposide response by multiplying the p53-driven production rates of MDM2, p21 and BTG2 with *α* for the etoposide data. Moreover, we made the stress decay rate compound-specific. The equations for the resulting p53 signalling model are:

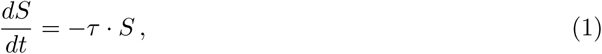

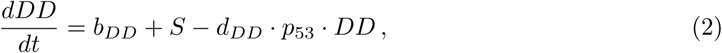

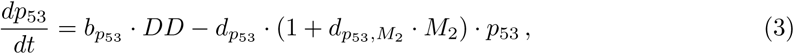

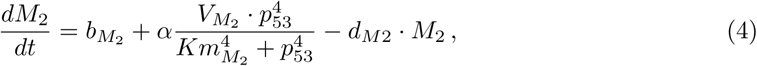

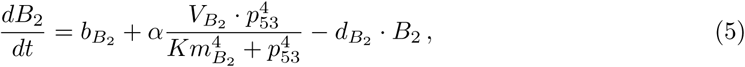

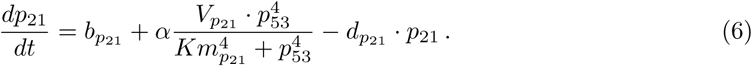

The description of each parameter and their values can be found in Supplementary Table 1. To ensure the model would be in steady state when simulating without stress, we used an approach similar to [32]. Specifically, we defined the degradation rates in each equation by a steady-state constraint based on the other parameters in that equation. This was achieved by setting the ODE equal to 0, and then solving the equation for the selected degradation rate considering absence of molecular stress (*S* = 0) and the other variables set to their initial state values.

#### Cell cycle models with mitosis

To mathematically describe cell cycle progression as influenced by DDR signalling, we built a simplified cell cycle model and integrated the p21 and BTG2 state variables. In the cell cycle model, we distinguished four cell cycle phases, i.e., G0, early G1, G1 and S-G2-M. At the start of the cycle, cells are in early G1 phase. Each cell can either progress in the cell cycle and enter G1 phase with rate *r_G_*_1_ or exit the cell cycle into quiescence (G0) with rate *r_exit_*. From G0, cells can re-enter the cell cycle in G1 phase with rate *r_entry_*. Cells in G1 progress to S-G2-M phase with rate *r_SG_*_2*M*_ and subsequently divide with mitosis rate *r_mit_*. Mitosis of a cell leads to the birth of two daughter cells that are in early G1 phase. To explain the accumulation of G1 cells in control conditions, we included resources *R* as additional state variable. Resources, such as growth factors, nutrients and unoccupied space, stimulate the transition from G1 to S-G2-M, but are depleted with a rate depending on the total number of cells scaled with factor *s*. Modelling the effect of p21 and BTG2 on cell cycle progression required integration of these proteins in the cell cycle model. Because p21 and BTG2 are both known to play a role in G1 and G2 arrest, we made the transition rates from G1 to S-G2-M and from S-G2-M to early G1 phase dependent on these proteins. We achieved this by dividing the appropriate transition rates by the scaled amounts of p21 and BTG2, i.e., *P*_1_ and *P*_2_, where

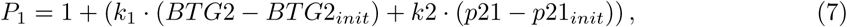

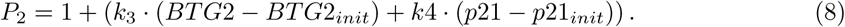

Note that in these equations we considered inhibition of cell cycle progression only when BTG2 and p21 expression exceeded their background levels, i.e., any such inhibition in control conditions is accounted for within the regular phase transition rates. We achieved this by deducting the initial values for BTG2 (*BTG*2*_init_*) and p21 (*p*21*_init_*) from their simulated values. The parameters *k*_1_, *k*_2_, *k*_3_ and *k*_4_ scale the relative influence of p21 and BTG2 on the cell cycle transitions. The ODEs for the cell cycle model thus became:

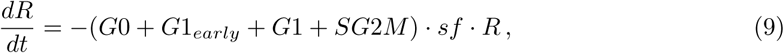

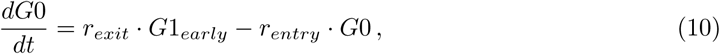

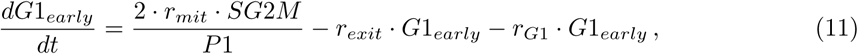

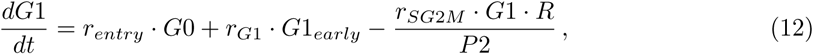

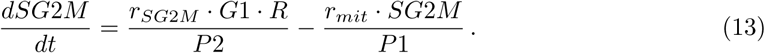

The total number of cells is not defined by a separate ODE but by summing the cells in all four phases. Initially, we calibrated the transition rate parameters to the data without drug exposure, followed by calibrating *k*_1_ to *k*_4_ with the cisplatin cell cycle data. We made changes to this default model (M1) to explore the effect of different model components on the simulations. Specifically, to examine the influence of resources, we removed the dependency of the G1 to S-G2-M transition on *R* and recalibrated all parameters (M2). Model M2 performed worse so for all other models we used the base parameters of model M1. To investigate the contribution of p21 and BTG2 to cell cycle progression, we set the *k_i_* parameters one by one to 0 while recalibrating the others (leading to models M3, M4, M5 and M6).

#### Cell cycle models with mitosis and endoreplication

Because cells exposed to DNA-damaging drugs can endoreplicate [22, 33], we included this process in our models. Specifically, we defined model M7 by replacing Equation 12 by:

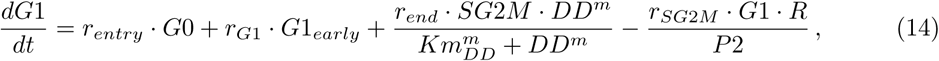

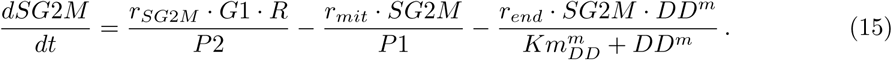

In the additional term for endoreplication, *r_end_* is the maximal rate at which cells can endoreplicate, *Km_DD_* is the amount of DNA damage for which the rate will be half maximal and *m* is a Hill coefficient.

For model M8, we tested whether endoreplicating cells could also first transition to early G1 phase instead of directly to G1 (i.e. first become colourless instead of red). To this end we used the same parameters as in model M7 but Equation 14 was changed back to Equation 12 and Equation 11 was changed to

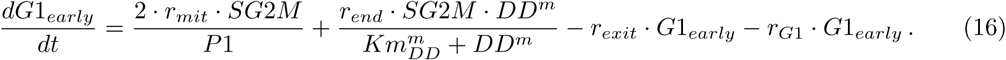

Model M8 fit the cell cycle data worse than model M7, so we did not pursue further investigation with this model formulation.

We used the best fit for model M7 to examine whether the same model could fit the data for etoposide exposure. Additionally, we attempted recalibration of the parameters *k*_1_-*k*_4_, *r_end_*, *Km_DD_* and *m* (model M9) or parameters *r_end_*, *Km_DD_*, *m* and *r_mit_* (model M10) for the etoposide data.

For model M7 for cisplatin and M10 for etoposide we determined the fraction of cells that was predicted to endoreplicate or divide based on these models. Specifically, for each time step of 0.1 hours, we determined the amount of cells leaving the S-G2-M phase by mitosis (given by the term 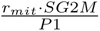) or by endoreplication (given by the term 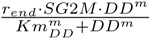). All these values were combined to determine the percentage of mitosis versus endoreplication over the whole timespan for each concentration.

#### Model parameter calibration

Parameter calibration of all models was done as described in [29]. In short, we used the sensitivity equations [34] to efficiently determine the path of steepest descent towards a local optimum. We performed optimisation of the objective function with the least squares method of the SciPy package in Python version 3.6.7. To find the global optimum in the parameter space, we initialised the models 50 times with a parameter set retrieved by systematic sampling of the parameter space with Latin hypercube sampling [35]. The model fitting results with the lowest cost were considered the global optima. The parameter values used to simulate all models can be found in Supplementary Table 1-4.

#### Data and code availability

All data and code to run the models and replicate the figures will be made available in a public repository upon publication.

## Results

### HepG2 cells transiently arrest in G2 phase after cisplatin exposure

To study the effect of DNA damage on the cell cycle, we generated HepG2-FUCCI cells. These cells contain red fluorescent protein-labelled Cdt1 (Cdt1-RFP) and green fluorescent protein-labelled Geminin (Geminin-GFP) and therefore display a red or green colour depending on the expression of Cdt1 and Geminin (Figure 1A and B). During G1 phase, Cdt1 protein levels increase, whereas Geminin remains absent, which manifests as red coloured cells. Cdt1 is degraded under the influence of Geminin, which builds up during the transition from G1 to S phase. The presence of both Cdt1 and Geminin therefore briefly yields cells with a yellow colour. In S, G2 and M phase, cells colour green due to high Geminin expression. However, after mitosis, Geminin is quickly degraded which renders the cells colourless. We measured the intensity of the RFP and GFP signals in HepG2-FUCCI cells with confocal microscopy (Figure 1B) to obtain the distribution of cells per cell cycle phase over time. Cells were either left untreated, i.e., in the presence of DMEM, or exposed to cisplatin just before imaging. As expected, because we did not experimentally synchronise the cells, they were in different phases at the start of imaging (Figure 1C, leftmost images). Most cells were colourless (i.e., no red or green signal) at 0 h and only gained colour over time, after entering the cell cycle (Figure 1C). In addition, the images clearly showed that cell cycle progression is influenced by the exposure to cisplatin.

**Figure 1:**
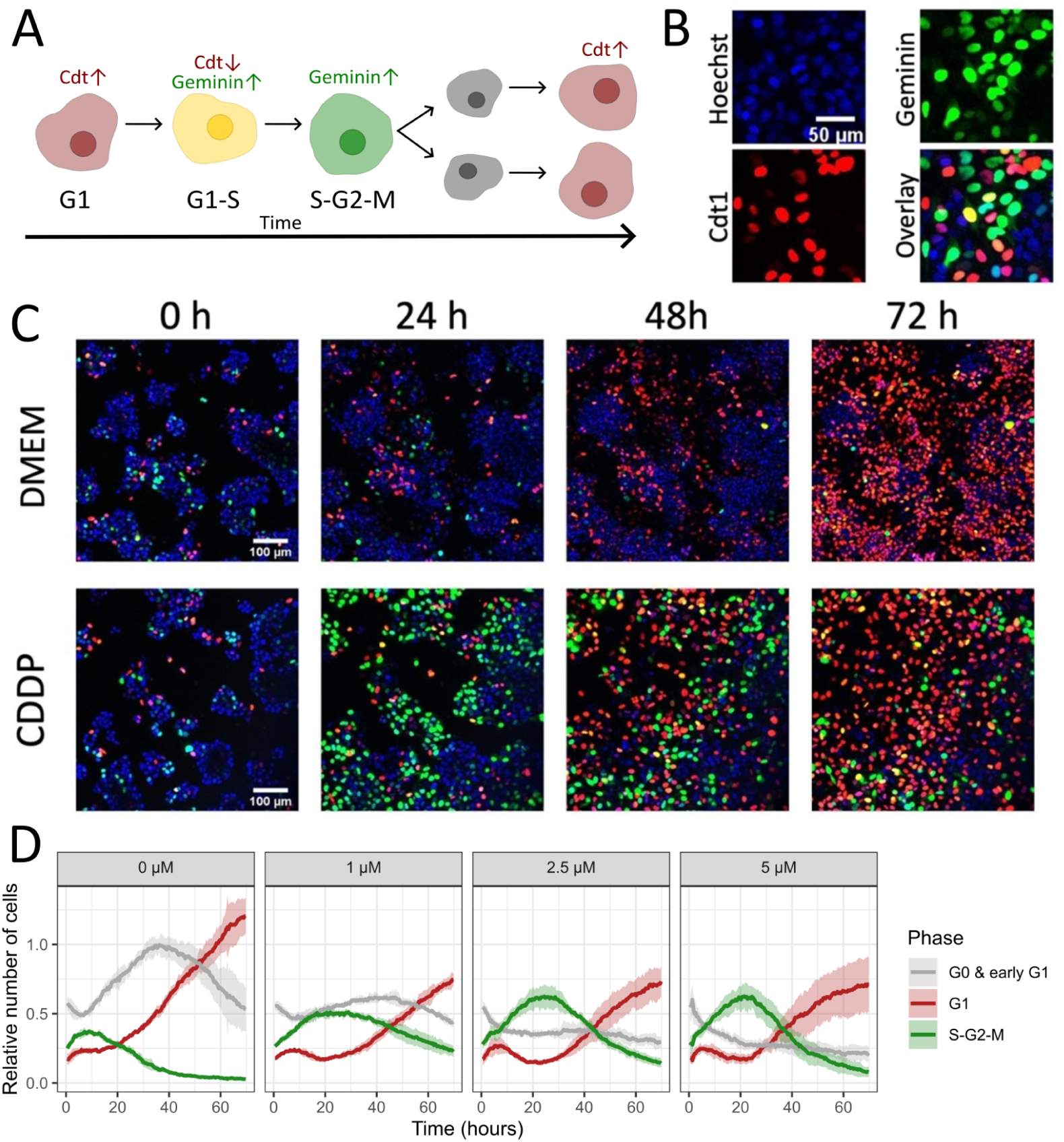
Cisplatin induces transient G2 cell cycle arrest. (A) Schematic overview of fluorescent signals during cell cycle progression in HepG2-FUCCI cells. Cells are red if Cdt1-RFP is expressed and green if Geminin-GFP is expressed. Yellow cells with both Cdt1 and Geminin expression are in a transition phase between G1 and S. Right after mitosis, cells are colourless (indicated with grey). (B) Example confocal microscopy images of HepG2 nuclei with Hoechst (blue), Cdt1 (red) and Geminin (green) fluorescent signals and their overlay. If both Cdt1 and Geminin are expressed, nuclei have a yellow colour. (C) Example images of fluorescence in HepG2-FUCCI cells in control, medium (DMEM) condition and after 5 µM cisplatin (CDDP) exposure. Scale bar is 100 µm. (D) Dynamics of the mean number of cells (solid lines) and standard deviation (shaded areas) in early G1 and G0, G1, and S-G2-M cell cycle phases at different cisplatin exposure concentrations.

To determine the cell cycle phase per cell at every measurement time point, we used the Cdt1-RFP and Geminin-GFP expression intensity. We noticed that cells with a high RFP intensity always transmitted some GFP signal as well, whereas cells with a high GFP intensity did not necessarily also transmit an RFP signal (Supplementary Figure 2A). Because there is some overlap between the emission spectra of GFP and RFP, the co-occurrence of GFP in RFP-positive cells was likely due to bleed-through. We corrected for this minor effect by applying a scaling factor to the GFP intensity based on the Cdt1 intensity (Supplementary Figure 2B). Based on the RFP and corrected GFP intensities, we determined a threshold below which cells were considered colourless (i.e., no RFP or GFP expression). Because most cells had visually very low RFP and GFP expression at the start of imaging (Figure 1C and Supplementary Figure 2C), we chose the threshold such that most cells were classified as colourless at 0 h. These cells were presumably residing in a quiescent G0 phase at 0 h and entered the cell cycle during imaging, or these cells may have just undergone mitosis and therefore were in the colourless early G1 phase. Cells with a Cdt1 expression higher than the threshold were considered to reside in G1 phase, whereas cells with a Cdt1-RFP expression lower but a Geminin-GFP expression higher than the threshold were classified as S-G2-M phase cells (Supplementary Figure 2D). Based on this classification scheme, we estimated the number of cells in each cell cycle phase over time (Figure 1D). This analysis showed that in control conditions HepG2 cells rapidly arrested in G1, in most cases presumably after one cell division. In contrast, cells that were exposed to cisplatin first went through a temporary G2 arrest, indicated by the accumulation of cells in S-G2-M phase around 24 h, prior to a halt in G1 phase at 72 h. This effect was stronger for high than for low cisplatin concentrations, which was also illustrated by the slow cell population growth at high concentrations (Supplementary Figure 1A). In conclusion, cisplatin-induced DNA damage leads to a transient G2 arrest in HepG2 cells.

### Limitation of resources drives G1 arrest in control conditions

Because p21 and BTG2 are both known to play a role in cell cycle arrest, we examined the relative contribution of these proteins to changes in the cell cycle. For this purpose, we slightly adapted our previously established ODE model of the p53 response [30] (see Figure 2A and Methods). In this model, chemically-induced DNA damage leads to phosphorylation of p53. Active, phosphorylated p53 transcriptionally activates p21 and BTG2 in addition to MDM2, a protein that negatively regulates p53. Simulation of this model shows the dependency of MDM2, p21 and BTG2 expression on p53 activation (Supplementary Figure 3A). We expanded the model with a cell cycle model that simulates the number of cells in early G1, G0, G1 and S-G2-M phases with parameters that describe the transition rates between phases. The regulatory proteins p21 and BTG2 were considered to jointly inhibit the transitions from G1 to S-G2-M phase and from S-G2-M to early G1 (Figure 2A). In addition, we included a fifth state variable, called ‘resources’ in the model that described the availability of resources such as growth factors, nutrients and free space, and that also affected the transition from G1 to S-G2-M phase.

**Figure 2:**
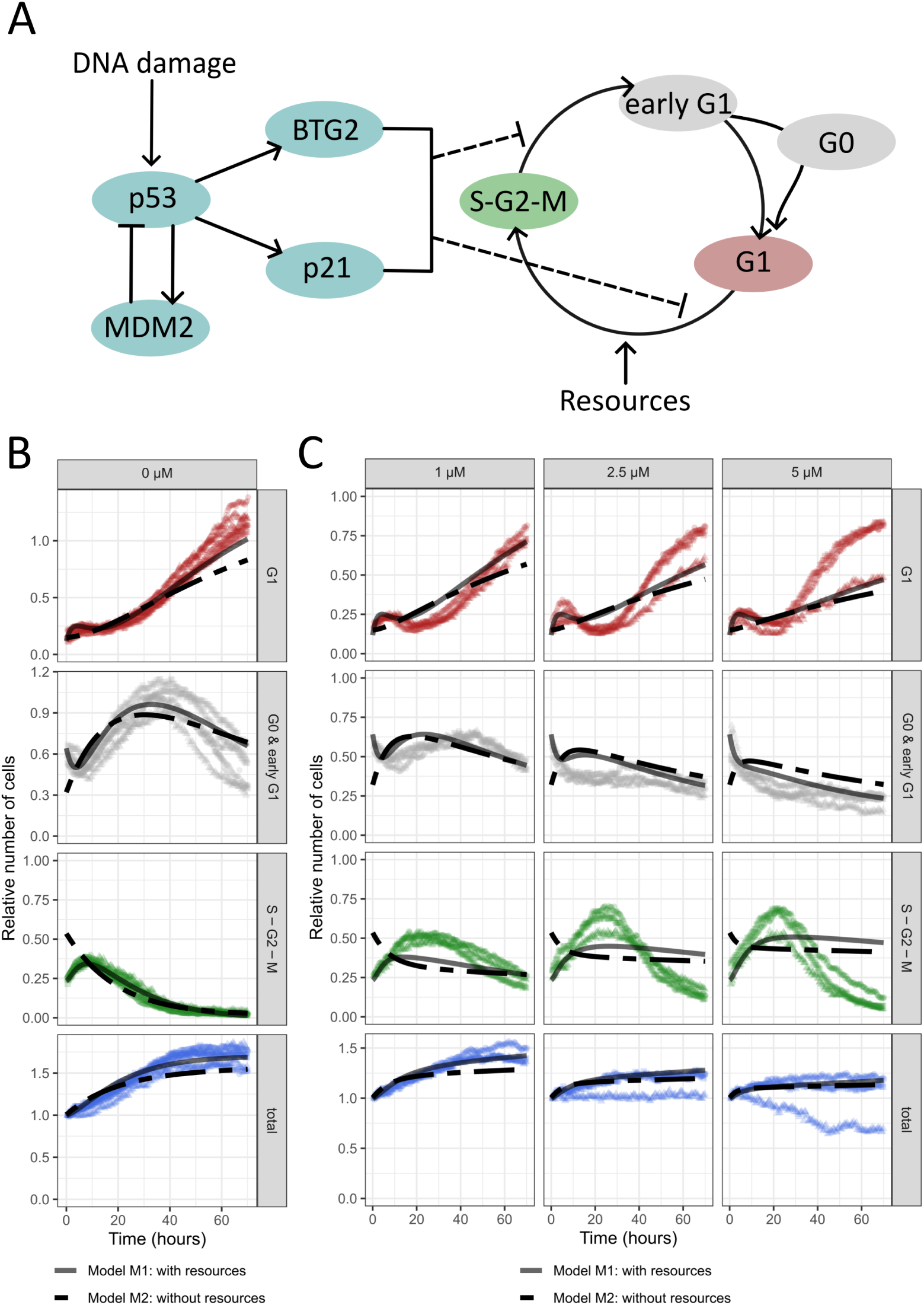
Depletion of resources leads to G1 arrest. (A) Illustration of the structure of our ODE model, with p21 and BTG2 that can inhibit cell cycle progression in both G1 and in S-G2-M phase. (B-C) Model simulations (M1, solid lines; M2, striped lines) and the normalised experimental data in control conditions (B) and after cisplatin exposure (C). Data points of different colours represent the three independent biological replicates. Model M1 includes depleting resources and model M2 excludes effects of resources.

Model parameter calibration to the experimental data generated a proper fit in control conditions (Figure 2B). Specifically, in the optimal model solution, G0 cells exited the G0 phase at the start of the simulation and entered the cell cycle (Supplementary Figure 4). Most cells remained arrested in G1 phase directly after entering the cell cycle or after one cell division, indicated by the stalling of cell population increase around 50 h at a cell count of approximately 1.7 times larger than its initial count (Figure 2B, bottom panel). The model predicted that resources were largely depleted around 30 h (Supplementary Figure 4). Importantly, omission of the effect of resources led to a worse fit, both in control conditions and after cisplatin exposure (Figure 2B and C). Without resources, the accumulation of cells in G1 phase was underestimated (Figure 2B, top panel) and there was a qualitative mismatch for the S-G2-M (green) and G0 and early G1 (colourless) cells at early time points. Specifically, the simulation without resources exhibited an initial increase in G0 and early G1 cells and a decrease in S-G2-M cells, which was opposite in the experimental data. Therefore, the model with resources is more suitable to describe the cell cycle dynamics, indicating that exhaustion of nutrients, growth factors or local space availability renders a plausible explanation for the observed G1 arrest in the control situation.

### Modelling predicts that endoreplication occurs upon cisplatin exposure

The experimental data from the cisplatin exposure conditions could be reasonably described with our model, except for the decrease in S-G2-M phase cells at late time points (Figure 2C). We examined the estimates of the parameters connecting BTG2 and p21 to the cell cycle to investigate whether we could pinpoint which links exhibited the strongest regulation. This analysis suggested that only p21 influenced the G1 to S-G2-M transition (see Supplementary Table 3, model M1). However, simulation results were similar when we made the inhibition of the G1 to S-G2-M transition solely dependent on BTG2 by fixing k4 to zero (model M3, Supplementary Figure 5A) or when we completely removed the dependency of the G1 to S-G2-M transition on the proteins (model M4, Figure 3A). This suggests that also in the presence of cisplatin the limited resources are the main driver for the G1 arrest at 72 h.

**Figure 3:**
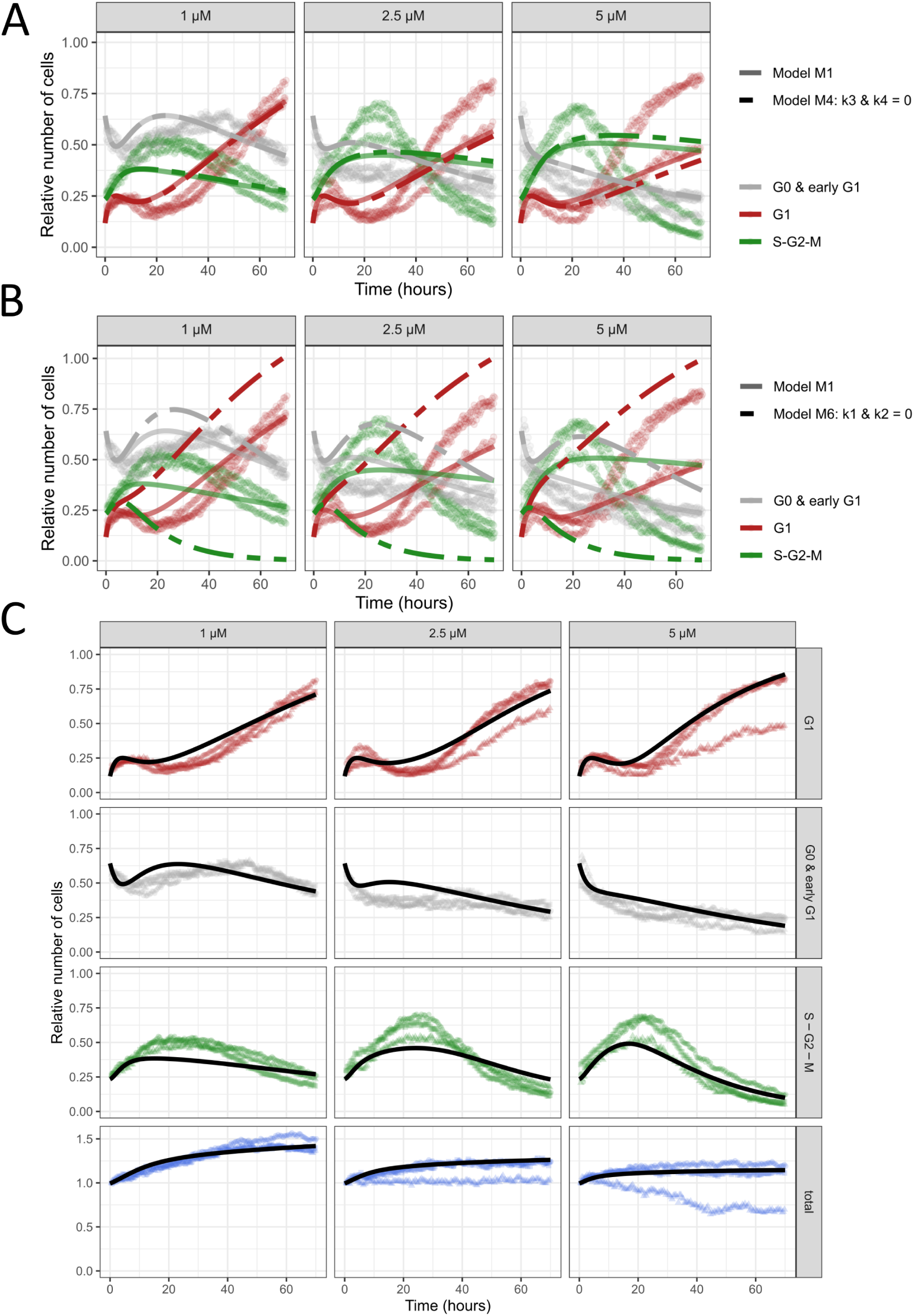
Endoreplication is required to describe decrease in S-G2-M arrest at late time points. A&B) Comparison of model with all inhibitory links from p21 and BTG2 to G1 and S-G2-M checkpoints present (model M1, solid lines) and model without inhibition of the G1 to S-G2-M transition by p21 and BTG2 (model M4, striped lines in (A)) or without inhibition of the S-G2-M to early G1 transition (including mitosis) by p21 and BTG2 (model M6, striped lines in B). C) Simulations of model with endoreplication (model M7, solid lines). Symbols of different colours represent the three biological replicates.

Further inspection of the estimated parameter values suggested that BTG2 was responsible for the S-G2-M phase arrest upon cisplatin exposure (Supplementary Table 3, model M1). However, if we made the S-G2-M to G1 early transition solely dependent on p21 by fixing k1 to zero (model M5, Supplementary Figure 5B), the fit was still similar. Nevertheless, either *k*1 or *k*2 is required to be non-zero because setting both parameters equal to zero leads to very poor predictions (model M6, Figure 3B): In that case the simulations lack the arrest in S-G2-M phase and overpredict the number of cells in G1 phase. Thus, BTG2 and/or p21 are drivers for the S-G2-M arrest in cells exposed to cisplatin.

The experimental data shows that cisplatin-exposed cells continued cycling after the transient S-G2-M arrest, yet the model simulations predicted a longer arrest (Figure 3A) due to the continued high expression of BTG2 or p21 until 72 h (Supplementary Figure 3A). This suggests that other factors are forcing cell cycle progression, irrespective of continued high BTG2 or p21 expression that inhibits this progression. One potential explanation is that cisplatin exposure can cause cells to endoreplicate [22, 36], i.e. cells alternate S and G phases without mitosis [19] and thus become polyploid. To incorporate this process into our model, we included a mathematical term to transition a fraction of S-G2-M cells directly to G1 phase without mitosis (see Methods). We considered the fraction of cells skipping mitosis to depend on the amount of DNA damage (Supplementary Figure 4), as this is a driver for endoreplication [18]. Calibration of this model (M7) resulted in simulations that properly capture the number of cells in S-G2-M phase, also at late time points (Figure 3C). One might wonder if the cells should not enter early G1 phase instead of G1 phase after skipping mitosis. To test this, we used the same parameters as in model M7 but with endoreplicating cells transitioning to early G1 phase instead of G1 (model M8, see Methods). This resulted in an increase in early G1 cells that did not match the data for the highest cisplatin concentration (Supplementary Figure 6). In conclusion, our model results suggest that cisplatin-exposed cells endoreplicate and that they swiftly transition to G1 phase when they skip mitosis instead of first becoming colourless as after mitosis.

### Modelling predicts massive polyploidisation upon etoposide exposure

To investigate the generality of our findings that S-G2-M arrest by p21 or BTG2 and endoreplication are both key processes driving cell cycle progression, we next aimed to test whether the resulting model could be applied to another DNA-damaging compound: etoposide. This is a DNA topoisomerase II inhibitor, which prevents ligation of DNA double-strand breaks during S and G2 phase [37]. As etoposide is a topoisomerase II inhibitor, it is well-known to induce endoreplication [33, 21, 18]. Exposing HepG2 cells to etoposide also leads to p21 and BTG2 induction (Supplementary Figure 3B). Therefore, we used the same p53-response model as for cisplatin, which resulted in an overestimation of the p53 levels upon etoposide exposure (Supplementary Figure 3B, purple line). When we introduced a rescaling factor for the effect of p53 on its downstream targets, this model reasonably described the etoposide data (see Methods; Supplementary Figure 3B, green line).

Exposure of HepG2-FUCCI cells to low etoposide concentrations, i.e., 0.5 and 1 µM, caused a transient S-G2-M arrest (Figure 4A) as was the case for cisplatin (Figure 1D). In contrast to the S-G2-M arrest that became more prominent with increasing cisplatin concentrations, at high etoposide concentrations (i.e., 5 and 10 µM) cells quickly bypassed the G2-M checkpoint and continued to G1 phase, indicated by the rapid drop of cells in S-G2-M phase and increase of cells in G1 phase(Figure 4A).

**Figure 4:**
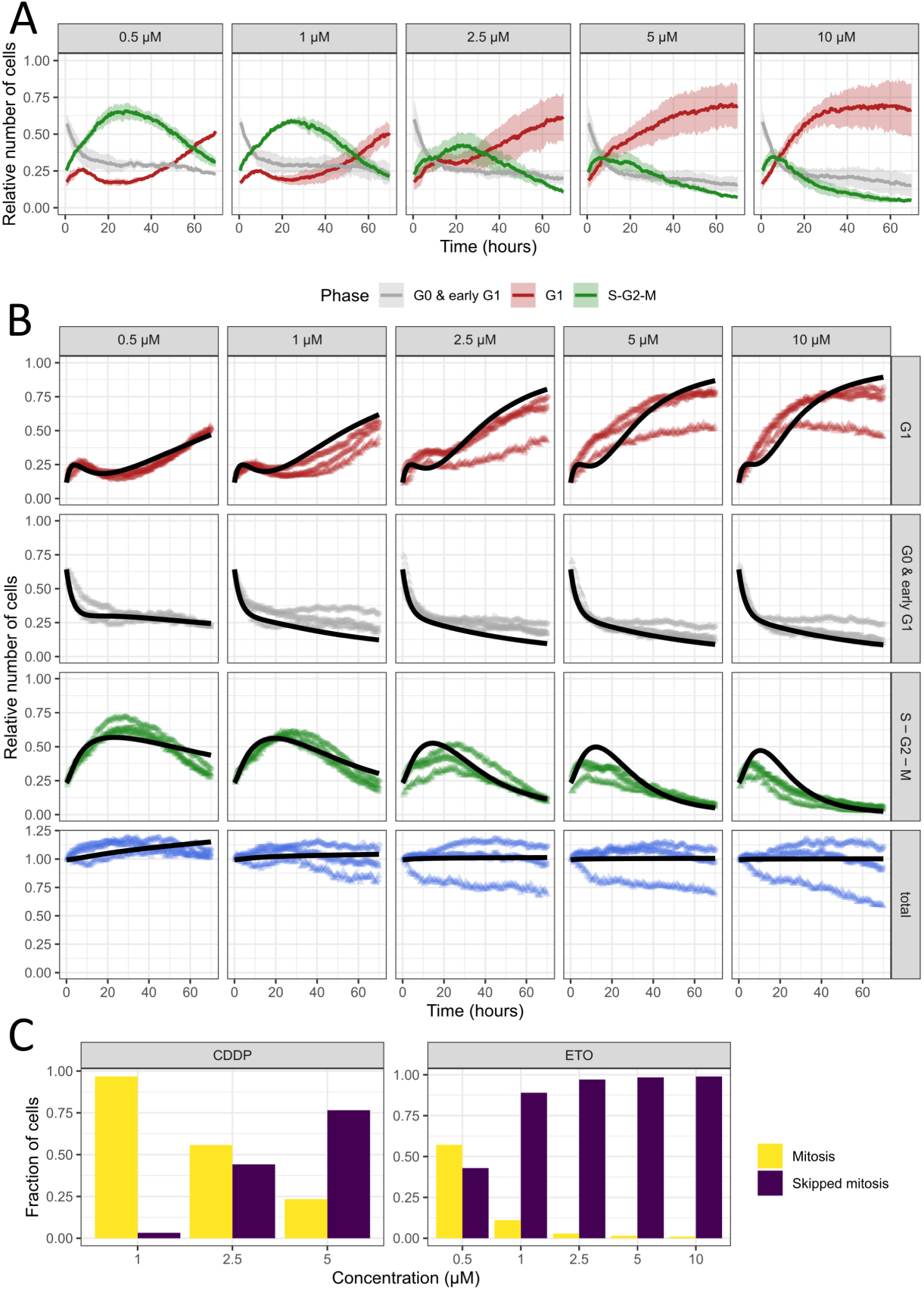
Model predicts massive polyploidisation for etoposide exposure. A) Dynamics of the mean number of HepG2 cells (solid lines) and standard deviation (shaded areas) in early G1 and G0, G1, and S-G2-M cell cycle phases at different etoposide exposure concentrations. B) Model simulations (M10, solid lines) after parameter estimation and the normalised experimental data following etoposide exposure. Symbols of different colours represent the three independent biological replicates. C) Predicted fraction of mitosis versus skipped mitosis events upon exposure to different concentrations of cisplatin or etoposide based on models M7 and M10 respectively.

To examine whether our model could describe these cell cycle progression data, we first simulated the model with endoreplication without changing any parameters from model M7 (Supplementary Figure 7A), i.e., the only difference with the cisplatin-based model lies in the input from the p53 signalling model (in the form of p21, BTG2, and DNA damage). With this model we obtained decent predictions for high concentrations but could not generate a good prediction of the observed S-G2-M cell cycle arrest at low etoposide concentrations. Furthermore, we noticed that proliferation was greatly impaired already at 0.5 µM etoposide, which was clear from the lack of growth of the cell population as a whole (Figure 4B, bottom panels) in combination with an absence of cell death (Supplementary Figure 1E and F). Because cells exposed to etoposide can arrest in G2 phase and skip mitosis already at low concentrations (found in NMuMG cells) [33], we expected that recalibrating the cisplatin-specific cell cycle parameters (*k*_1_, *k*_2_, *k*_3_, *k*_4_, *r_end_*, *Km_DD_* and *m*) would generate an improved simulation (model M9). Indeed, this parameter adaptation greatly improved the fit to the low-concentration etoposide data (Supplementary Figure 7B).

From inspection of the model M9 parameters (Supplementary Table 4), it was clear that both *k*_1_ and *k*_2_ were at their maximal value. This implied that the S-G2-M to G1 phase inhibition by p21 and BTG2 was estimated as very strong. However, the p21 and BTG2 expression at low etoposide concentrations was only slightly higher than the basal expression in control conditions (Supplementary Figure 3B) and low in comparison to cisplatin exposure, whereas the arrest in S-G2-M phase after etoposide exposure was more pronounced than in cisplatin-treated cells. Therefore, the rapid S-G2-M arrest that occurred in HepG2 cells exposed to low concentrations of etoposide is likely promoted by factors other than p21 and BTG2. Consistently, when we recalibrated the rate of mitosis *r_mit_* (in model M10) instead of the parameters *k*_1_-*k*_4_ (as in model M9), we obtained fits of similar quality (Figure 4B).

Based on the calibrated cell cycle models (M7 for cisplatin and M10 for etoposide), we computed the relative number of cells leaving the S-G2-M phase both with and without mitosis for each condition and time step (Supplementary Figure 8). We also computed the total fraction of cells during the whole timespan that skipped or underwent mitosis (Figure 4C). This analysis predicted that the fraction of cells skipping mitosis was concentration-dependent for cisplatin, whereas for etoposide a large fraction of cells was predicted to skip mitosis for all concentrations. Regardless, the relative number of cells skipping mitosis was expected to increase with the concentration for etoposide as well (Supplementary Figure 8).

### Time-lapse imaging confirms occurrence of polyploidisation following exposure to cisplatin and etoposide

To test our model predictions, we performed additional time-lapse imaging with the same set-up but at higher magnification. Due to the increased magnification, it was now possible to track individual cells manually with high confidence and identify whether cells skipped or underwent mitosis. As predicted by our model, in the untreated wells we observed many dividing cells (Figure 5A), while we observed many cells transitioning to red or colourless without division in the wells exposed to high drug concentrations (Figure 5B).

**Figure 5:**
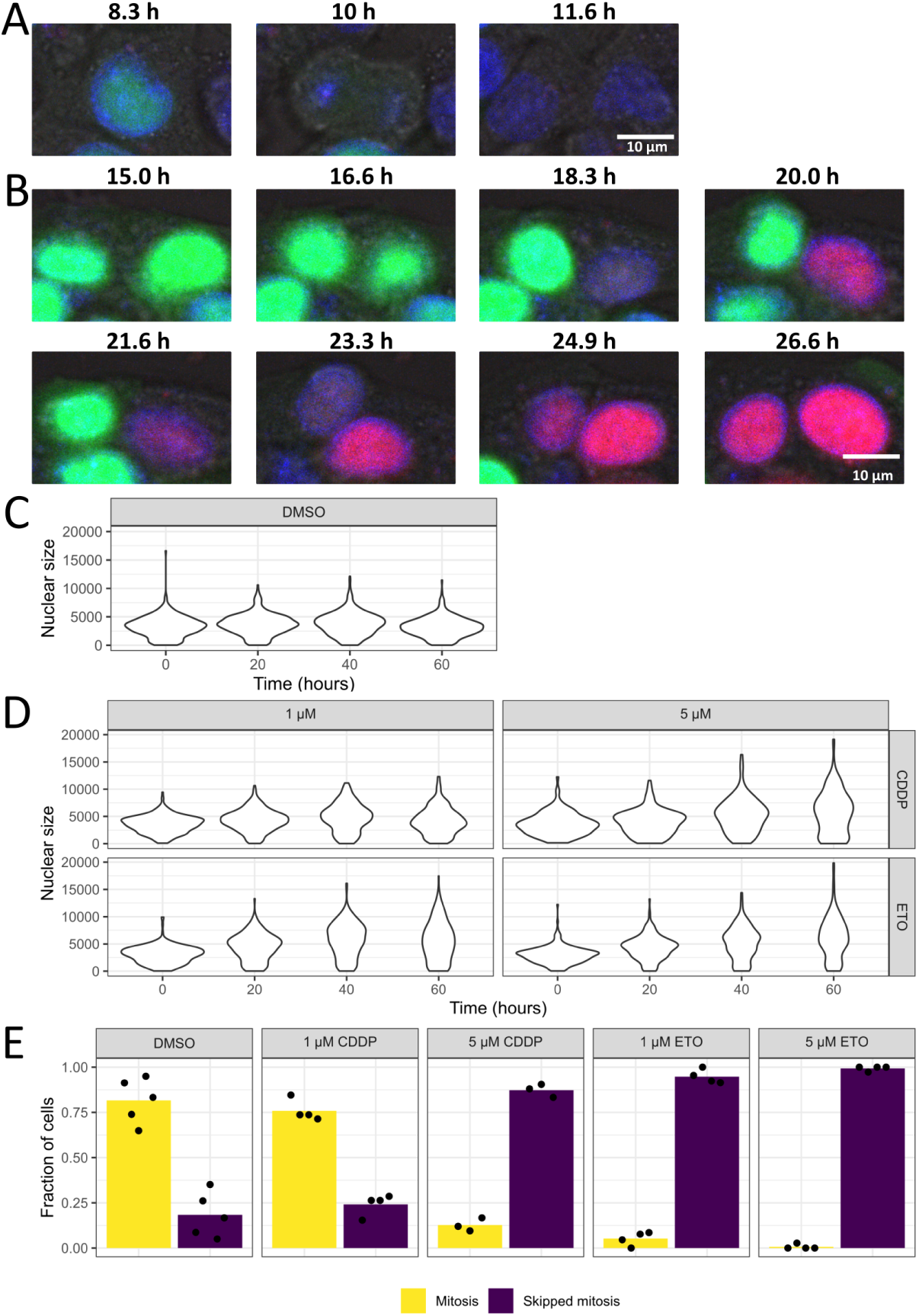
Imaging data support a large role of polyploidisation in cell cycle progression upon DNA damage. A) Example of a dividing cell in control conditions. At 8.3 hours the cell still expresses green fluorescence, at 10 hours the cell is in mitosis and at 11.6 hours the two daughter cells are visible without any red or green fluorescence. B) Example of two cells skipping mitosis in a well exposed to 5 µM etoposide. At 15 hours both cells are still bright green, the right cell transitions from green to red without a division between 15 and 20 hours. From 21.6 hours onwards, the left cell transitions to red without division in a similar manner. The scale bars in A and B are 10 µm. C & D) Distribution of nucleus sizes (in number of pixels) at 0, 20, 40 and 60 hours without exposure (C) or after exposure to 1 or 5 µM cisplatin (D, top row) or etoposide (D, bottom row). E) Quantification of cell-tracking, where we defined the fate of a green cell as mitosis if we could observe the division, and as skipped mitosis if the cell transitioned to red or colourless without division. Dots represent the fraction per biological replicate and the bars represent the mean fractions across replicates.

When cells endoreplicate, the amount of DNA increases, thus one would expect the nuclear size to increase as well. Therefore, following nuclear segmentation we quantified the size of each nucleus in the images. As expected, the distribution of nuclei sizes remained unchanged over time for the unexposed cells (Figure 5C). However, the distribution shifted towards larger nuclei in the exposed wells as time progressed (Figure 5D). For etoposide, we observed an increased number of large nuclei for both low and high concentrations, which matched our model prediction that the majority of cells become polyploid for most etoposide concentrations. For cisplatin, this increase was most profound for high concentrations and late timepoints. These results suggest that polyploidisation mostly occurred after exposure to etoposide or high cisplatin concentrations, which is in agreement with our model predictions.

We also performed manual cell-tracking to quantify how often cells skipped or underwent mitosis. We followed all green cells and counted an event as mitosis if we could observe a green nucleus dividing into two daughter nuclei. We classified an event as skipped mitosis if a cell would become colourless or red without a division (see Methods). We computed the fraction of cells skipping and undergoing mitosis based on the total number of classified cells. With this analysis, we found clearly higher fractions of cells that skipped mitosis (and thus lower fractions of dividing cells) for wells exposed to etoposide or a high concentration of cisplatin than for wells that were not exposed to a drug or to a low cisplatin concentration (Figure 5E). Furthermore, in wells exposed to a high concentration of the drugs, cells skipped mitosis early and often became red instead of colourless (Supplementary Figure 9). Both the early increase in cells skipping mitosis and increased amount of cells skipping mitosis for high concentrations is in accordance with our model predictions (Figure 4C and Supplementary Figure 8). In conclusion, these results demonstrate that polyploidisation is a key process during cell cycle progression of HepG2 cells exposed to DNA-damaging compounds.

## Discussion

Cell cycle progression is a complex cellular mechanism that involves numerous regulatory proteins. Externally induced cellular stress such as chemical-induced DNA damage can disrupt normal cell proliferation through the activation of proteins that interact with cyclins and CDKs, the drivers of cell cycle progression. Using HepG2 cells with an integrated FUCCI construct, we could follow cell cycle progression exposed to DNA-damaging agents cisplatin or etoposide. Clear temporal changes in the fraction of cells in G1 and G2 arrest occurred following exposure. With our models we showed that a combination of G1 arrest by limited resources, G2 arrest by BTG2 or p21, and DNA damage causing endoreplication could largely explain the observed cell cycle arrest as well as subsequent progression.

Cellular exposure to cisplatin and etoposide both caused a transient G2 arrest in HepG2-FUCCI cells. Our model could not distinguish between effects of BTG2 or p21 on the cisplatin-induced G2 arrest. Indeed, both p53 targets are reported to interfere with G2 arrest [14, 38], which indicates that either p21 or BTG2, or their combination could be responsible for the cisplatin-induced G2 arrest. In contrast, p21 and BTG2 expression barely increased at 0.5 µM etoposide exposure, whereas there was a profound transient accumulation of cells in S-G2-M phase. Our analysis showed that solely p21 and BTG2 expression was not sufficient to explain this etoposide-induced G2 arrest. Lowering the mitosis rate improved the fit, which suggests that other proteins than p21 and BTG2 could impact the phase transition rates. It is known that G2 arrest can occur in a p53-dependent and p53-independent manner [39, 40, 41, 42]. Moreover, whereas BTG2 can induce a G2 arrest [14], p21 seems mostly involved in the long-term maintenance of G2 arrest [38, 43]. Thus, perhaps the DNA damage sensor proteins ATM, ATR, Chk1 and Chk2 that activate p53 signalling elicited a p53-independent onset of G2 arrest [44, 45, 46] after 0.5 µM etoposide exposure, whereas the late and weak induction of p53-regulated protein p21 merely helped to sustain the G2 arrest. The possibility that different mechanisms establish the cisplatin- and etoposide-induced G2 arrests, indicates that the exact type of DNA damage influences the cellular response. In contrast to cisplatin that causes inter- and intrastrand crosslinks which activates p53 via ATR, Chk1 and Chk2 [47, 48, 49, 50], etoposide exposure generates double-strand breaks that lead to p53 activation via ATM and Chk2 [44, 22]. Although both compounds induce p53 activation and transient G2 arrest, the underlying molecular signalling cascade is different and therefore results in G2 arrest via different routes. With our work, we show that solely using the p53 downstream targets p21 and BTG2 to predict a DNA damage-induced G2 arrest is not always sufficient.

Incorporation of proteins upstream of p53 in experimental studies and the model will contribute to reliable predictions of cell cycle arrest.

Regardless of the sustained expression of p53 and its downstream targets at late time points, the cells continued to a G1 arrest after the G2 arrest in both exposure conditions. Cell cycle progression after G2 arrest was especially evident for cisplatin-treated cells and cells exposed to low (0.5 and 1 µM) etoposide concentrations. In contrast, high (5 and 10 µM) etoposide concentrations swiftly induced a G1 arrest. Because the number of cells did not increase after the S-G2-M to G1 transition yet cell death was minimal at these concentrations, we hypothesised that the cells must be endoreplicating. This same phenomenon has been previously observed in normal murine mammary gland (NMuMG) cells. NMuMG cells went into a transient G2 cell cycle arrest at 1 µM etoposide exposure, but after exposure to 10 µM etoposide they quickly endoreplicated and arrested in G1 phase [33]. Endoreplication also occurred in cisplatin-treated PROb colon carcinoma [36] and Chinese hamster ovary AA8 cells [51]. Although the observed accumulation of red HepG2 cells in our data could also mean that cells adapt and enter mitosis despite damaged DNA [52, 53, 1, 54], the total number of cells barely increased, pointing to a lack of proliferation. Consistently, modification of the model by including a separate mathematical term for a transition from G2 to G1 phase without mitosis strongly improved the model fit. Specifically, our model predicted that the majority of cells skip mitosis for 5 µM cisplatin and for 1 µM and higher concentrations of etoposide. By following the fate of all green cells in additional HepG2-FUCCI time-lapse imaging data of high magnification we confirmed these model predictions.

For our model, we considered all endoreplicating cells to transit immediately to G1 phase (red) instead of to G0 and early G1 phase (colourless) first. However, we observed that cells could also become colourless without division (Supplementary Figure 9). For high concentrations most cells transition immediately to G1 phase, but for lower concentrations, the cells transition to G0 and early G1 phase more often. A factor that may contribute to this striking difference involves nutrient and growth factor availability. For DMSO and low concentrations of cisplatin, the majority of cells divide, and this increases the number of cells, which is expected to lead to resource depletion. Consequently, such depletion could make cells arrest in G0 instead of committing to G1 phase. However, for low concentrations of etoposide there are hardly more cells than for high concentrations, yet generally cells skip mitosis later for 1 µM than for 5 µM etoposide (Supplementary Figure 9B). Moreover, some cells also become colourless at early timepoints or red at late timepoints. These observations suggest that additional factors besides resource availability influence the speed at which G1 is entered after a skipped mitosis. Thus, further research is required to investigate the cause of the differences in red and colourless cells for varying concentrations of DNA-damaging agents.

Our models for cisplatin showed that when endoreplicating cells would be considered to transition to early G1 instead of G1 phase, this would result in an increase in colourless cells for the highest cisplatin concentration, which was not observed in the data (Supplementary Figure 6, right column). However, for the lowest cisplatin concentration, the modelled and observed abundance of colourless cells would match reasonably well (Supplementary Figure 6, left column). This could be explained by the low number of cells endoreplicating at low cisplatin concentrations. Therefore, it would not be required to change this aspect of our model to represent the data. However, in a future application of the model, one could consider splitting the endoreplicating cells between cells transitioning to G1 early and cells directly transitioning to G1 phase in a concentration-dependent manner to better represent the biology.

In the time-lapse imaging data with high magnification we classified cells as skipping mitosis when they transitioned from green to red or colourless without division. Apart from endoreplication, there are other ways to transition from the S-G2-M phase to G1 phase without division. First, mitotic slippage is a process similar to endoreplication; in mitotic slippage, cells enter the M phase but do not finish it before continuing to the G1 phase, while endoreplication skips the M phase entirely [55]. Distinguishing between mitotic slippage or endoreplication would require additional imaging experiments with time steps of a few minutes to distinguish different mitotic phases [56]. Second, prior work in which NMuMG cells were exposed to 1 µM etoposide did not only endoreplicate yet many cells also underwent missegregation, which was characterised by chromosome fragmentation [33]. With our 20x magnification, it was impossible to distinguish this phenomenon, and we thus did not take it into account in our models. Third, bone cancer cells exposed to 20 µM etoposide were recently reported to rereplicate [57], which means that cells replicate their DNA multiple times within one S phase. If this is a generic phenomenon that also occurs in other cell types and for low etoposide concentrations, this could further amplify the polyploidisation in cells skipping the M-phase. In the future, it could be useful to develop a model that can describe various adverse cell fates, such as cell death, missegregation and rereplication.

We aimed to predict cell cycle behaviour based on the average p21 and BTG2 expression and estimated DNA damage in a population of cells. Although our HepG2 cells are genetically almost identical, the cells varied in the expression of proteins, and cell cycles were not synchronised. Such variability can exert their effect on the cell cycle of individual cells in various ways: differences in the time dynamics and extent of p53 expression determine whether a cell initiates cell cycle arrest or apoptosis [11]. Similarly, basal p21 expression controls quiescence or cell cycle entry [58], whereas p21 dynamics determine whether a cell remains in a proliferative state or becomes senescent [59]. In addition, cells respond differentially to genomic stress depending on the cell cycle stage [60]. Other factors such as the microenvironment [61, 62] and general stochasticity [63] can cause heterogeneity in the behaviour of individual cells. Thus, although a study into the relation between protein dynamics and cell fate on a population level can generate useful insights, single cell data can further elucidate the mechanistic regulation of cell cycle progression after genotoxic stress.

Integrating protein dynamics and cell cycle progression in one mathematical model is a useful approach to unravel the effect of low-level molecular interactions on a high-level process such as cell fate. We showed that p21 and BTG2 expression dynamics can largely predict (transient) cell cycle arrest following genotoxic stress. In addition to the inclusion of p53 downstream targets, extension of the model with the activation dynamics of kinases upstream from p53 could improve its predictive capacity. Furthermore, our models demonstrated a large role for polyploidisation in HepG2 cells exposed to DNA-damaging compounds, which we verified experimentally. Our findings can help to understand the quantitative mechanisms behind cell cycle regulation and predict the adverse effects of genotoxic chemicals in an early stage of drug safety testing.

## Acknowledgements

This project has received funding from the ZonMW InnoSysTox program under grant agreement no. 114027005 (SysBioToP-Moving). The authors gratefully acknowledge the imaging core facility, the Leiden Cell Observatory for their support and assistance in this work. The authors would like to thank the members of the SysBioTop-Moving consortium for scientific discussions related to this work.

## Supplementary material

### Supplementary Figures

**Supplementary Figure 1:**
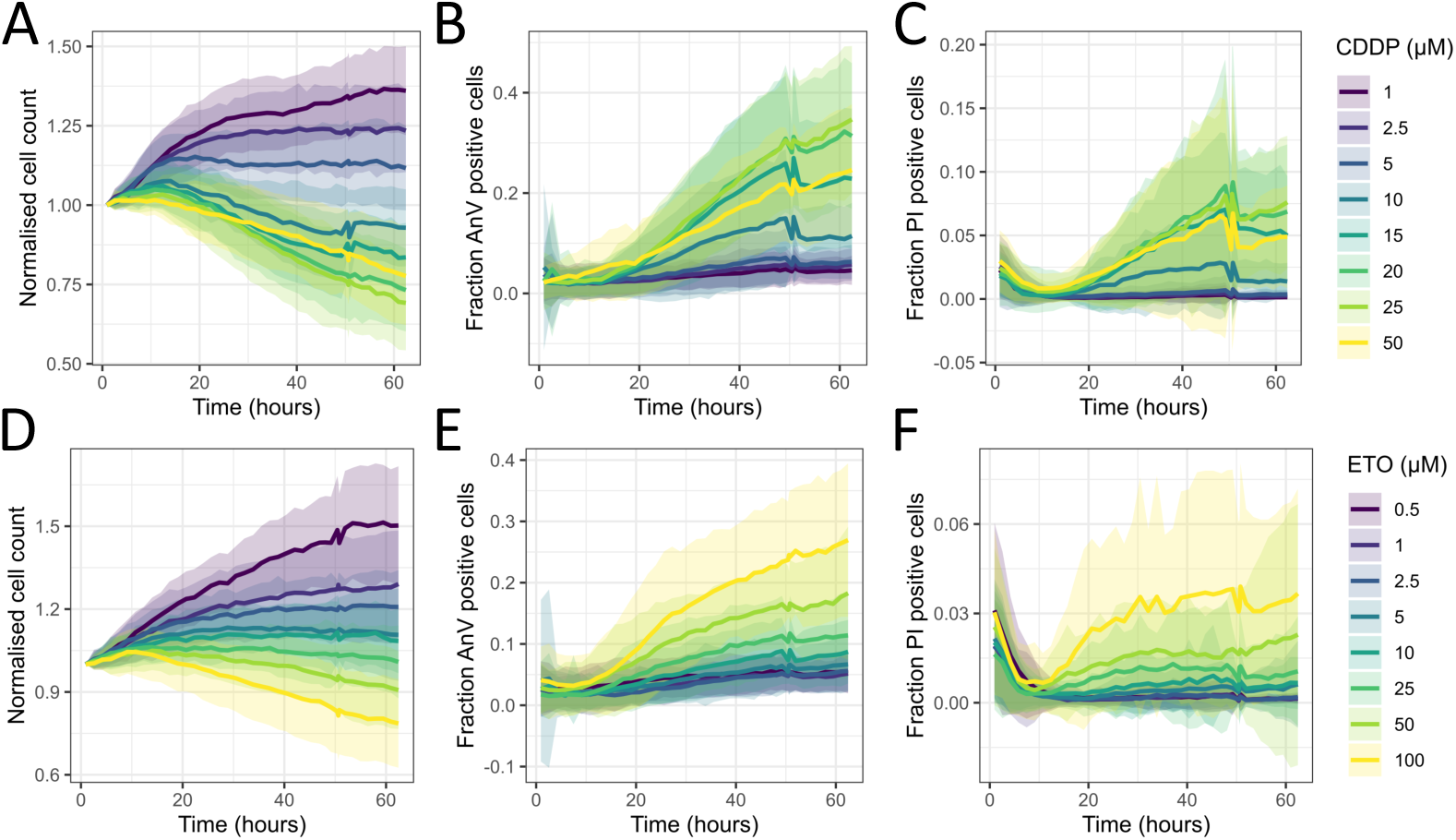
Viability data based on cell count, AnV and PI. Data for eight concentrations of cisplatin (A-C; CDDP) and etoposide (D-F; ETO). A&C) Normalised cell count, B&E) fraction AnV-positive cells and C&F) fraction PI-positive cells. The colour represents the concentration, the line is the mean, and the shaded area is the standard deviation across 31 biological replicates (at least three replicates for each of the 11 different types of reporter cell).

**Supplementary Figure 2:**
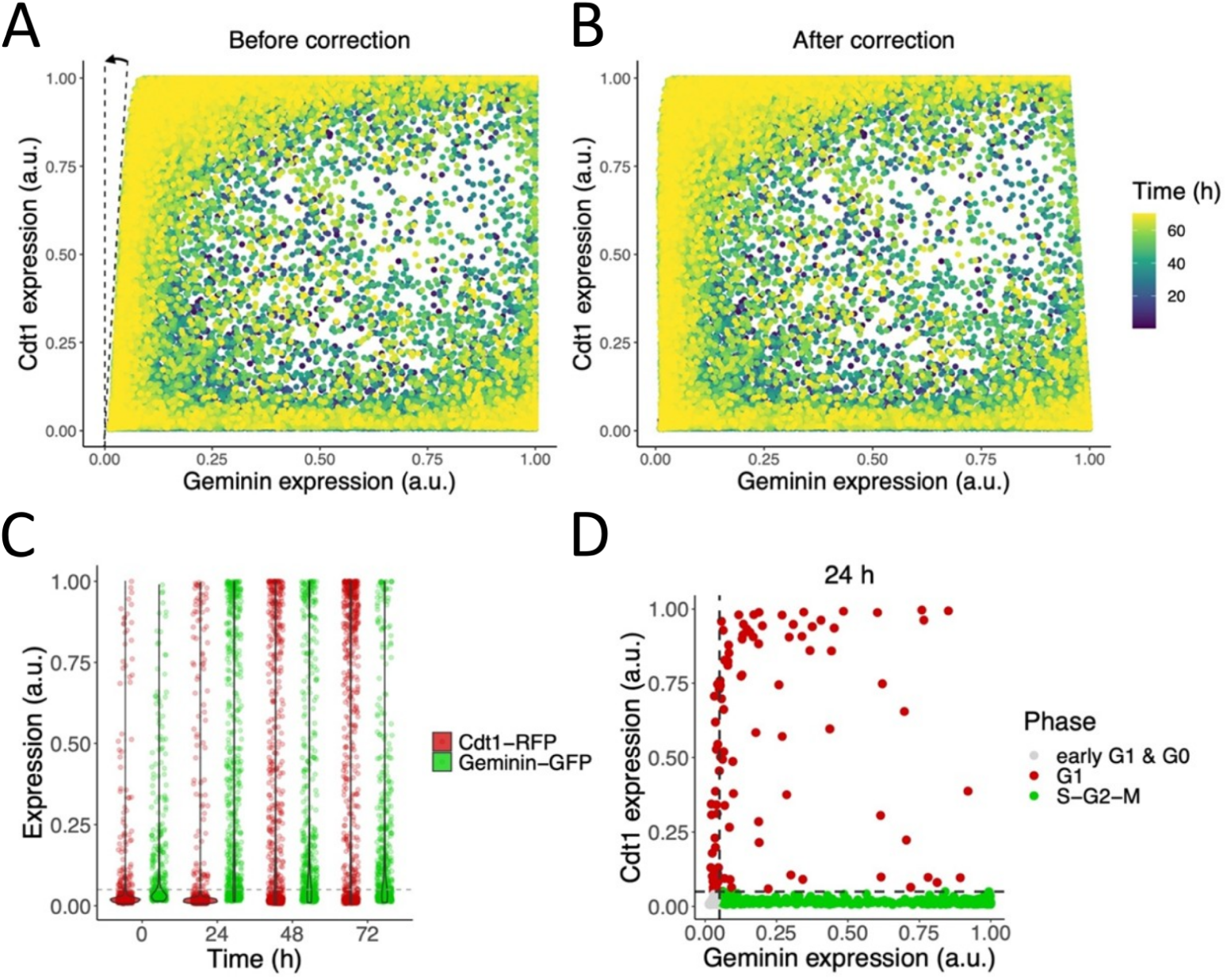
Quantification and classification of cell cycle phases based on Cdt1-RFP and Geminin-GFP intensities. (A-B) Example distribution of Cdt1 and Geminin expression in all cells from one image set, i.e., all time points at one position in a well, before (A) and after (B) spectral leakage correction. (C) Example distribution of Cdt1-RFP and Geminin-GFP intensities at 0, 24, 48 and 72 h after exposure in 5 µM cisplatin condition. The horizontal grey dashed line indicates the 5% threshold, below which Cdt1 and Geminin expression is low and classified as colourless. (D) Example image of Cdt1 and Geminin expression at 24 h after 5 µM cisplatin exposure. The dashed lines represent the utilised 5% expression thresholds, resulting in the classification of cell cycle phases as indicated by colours. In A-D each dot represents one cell.

**Supplementary Figure 3:**
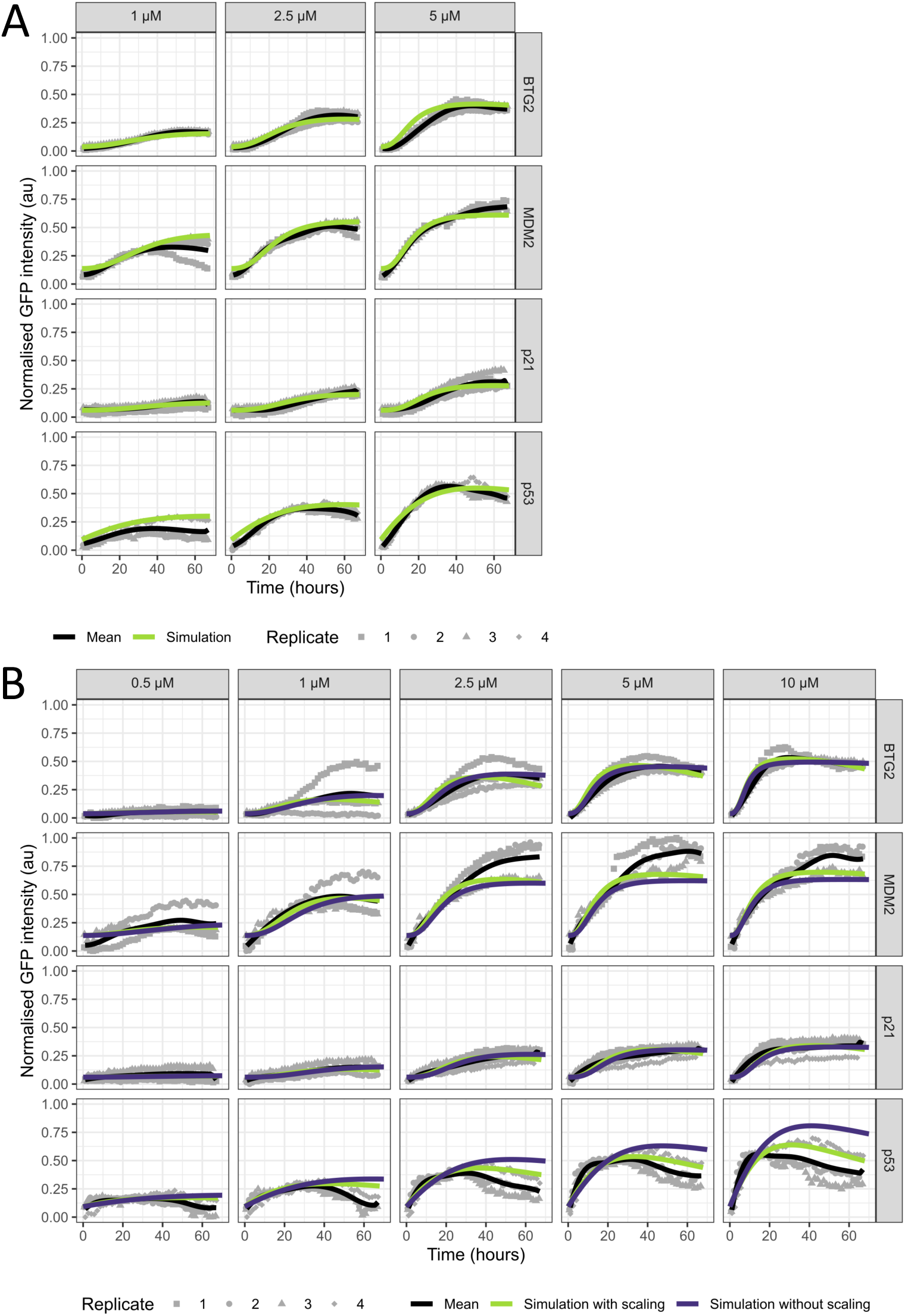
Experimental and simulated p53 signalling dynamics upon exposure to cisplatin or etoposide. Model simulations for cisplatin (A) and etoposide (B). For the etoposide simulations, scaling of the p53 target gene responses is either present (green line) or absent (blue line). The black solid line represents the mean and the grey symbols represent the three biological replicates.

**Supplementary Figure 4:**
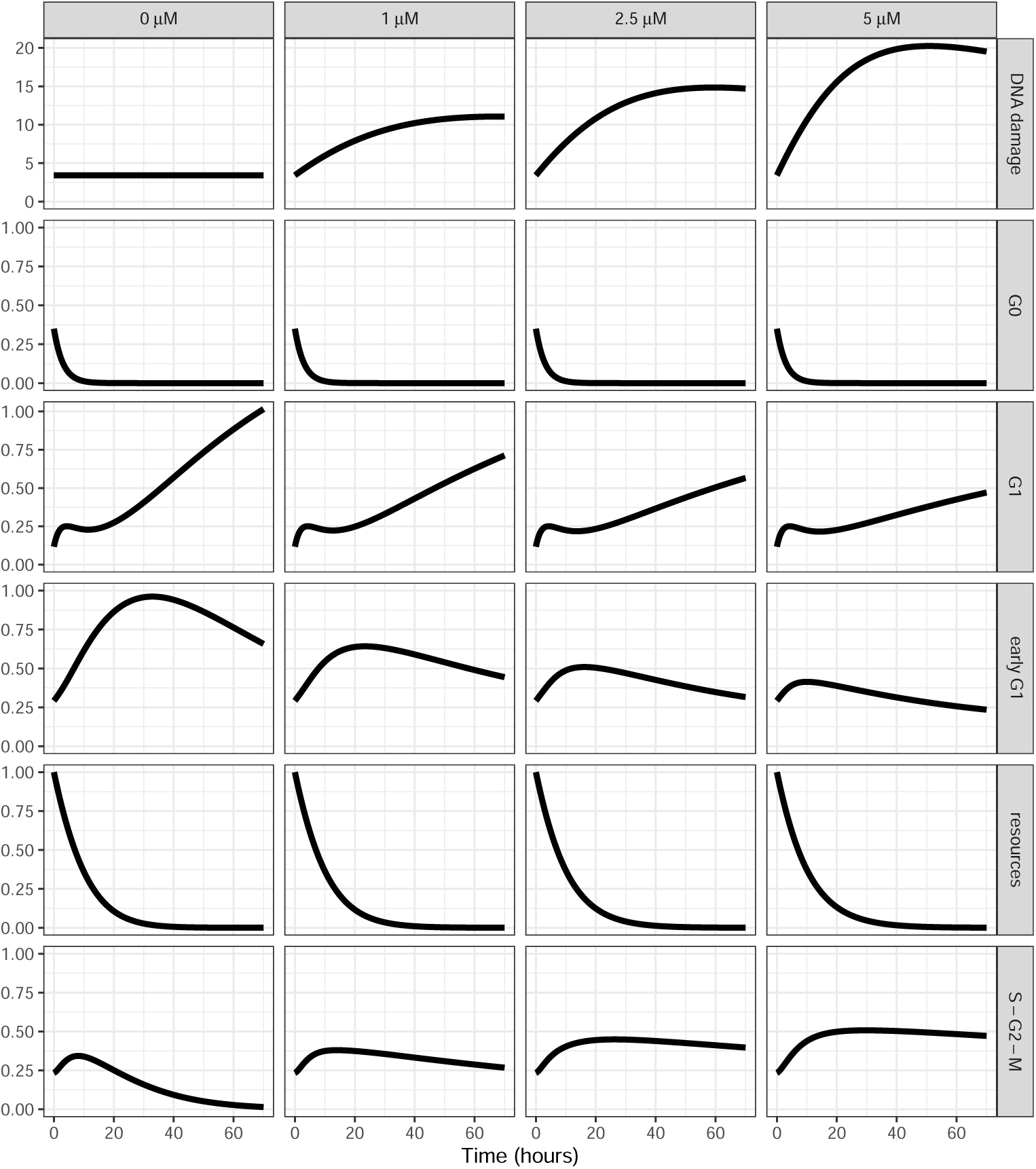
Inner states of the cell cycle model simulation after 0, 1, 2.5 and 5 µM cisplatin exposure. Simulation of the relative number of cells in the individual cell cycle phases and of the resources for model M1.

**Supplementary Figure 5:**
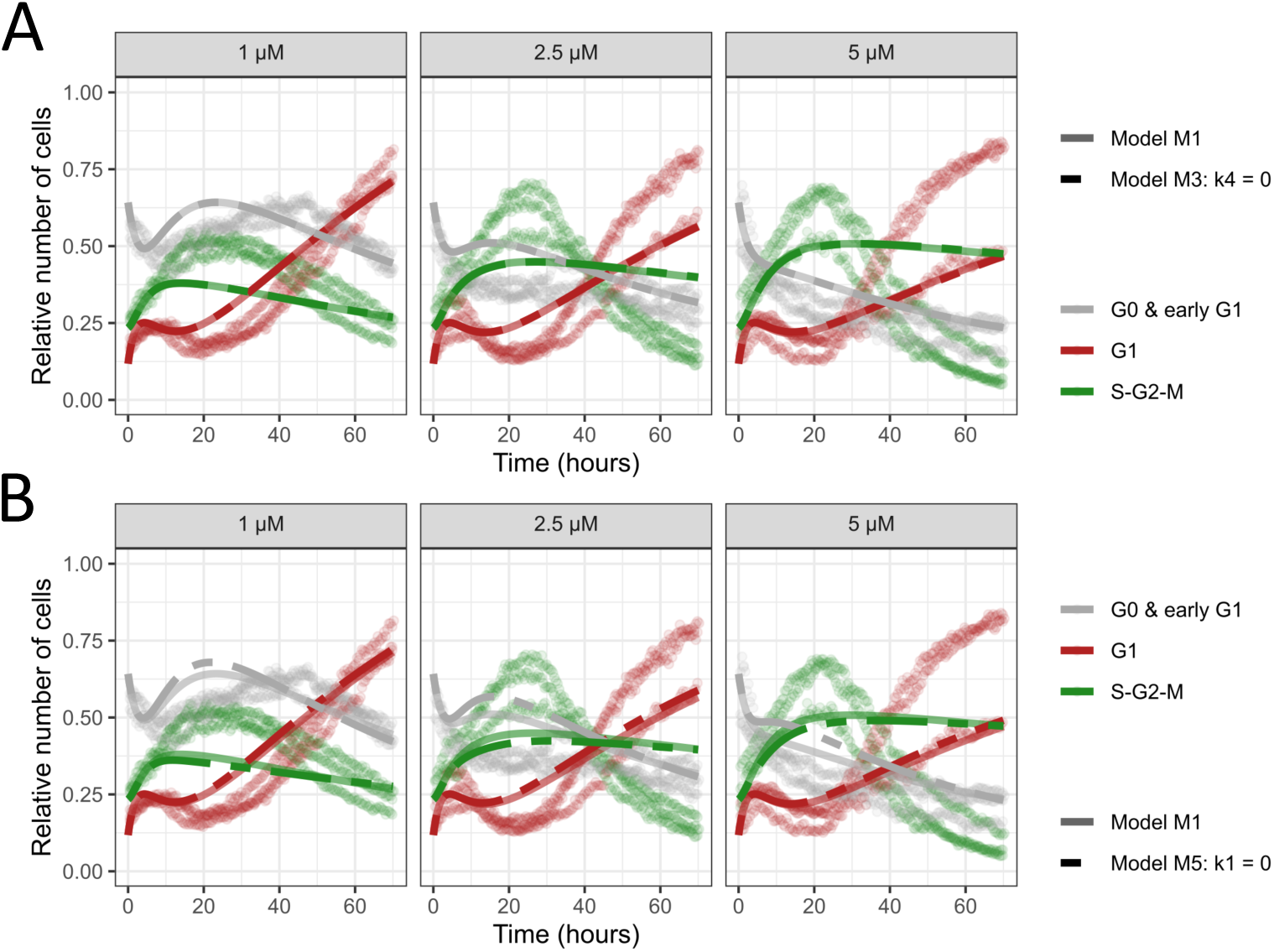
Effect of removing k4 or k1 from the model for cisplatin. A-B) Comparison of model M1 (solid lines) and model with inhibition of G1 to S-G2-M transition solely by BTG2 (model M4, striped lines in A) or with inhibition of mitosis solely by p21 (model M6, striped lines in B).

**Supplementary Figure 6:**
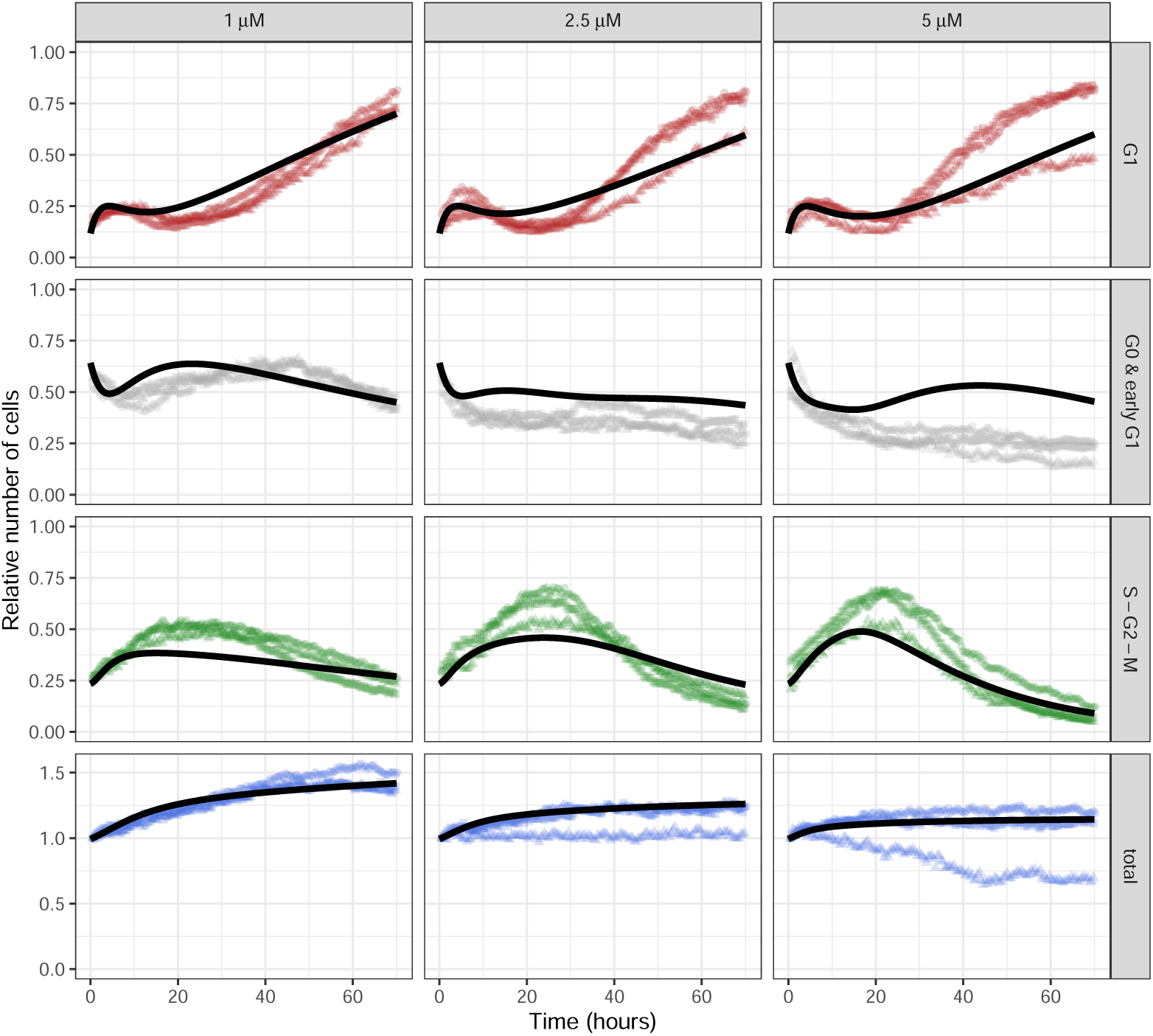
Effect of adding endoreplicating cells to early G1 rather than G1 phase. Simulations of model with endoreplicating cells added to early G1 phase (model M8, solid lines). Symbols of different colours represent the three biological replicates.

**Supplementary Figure 7:**
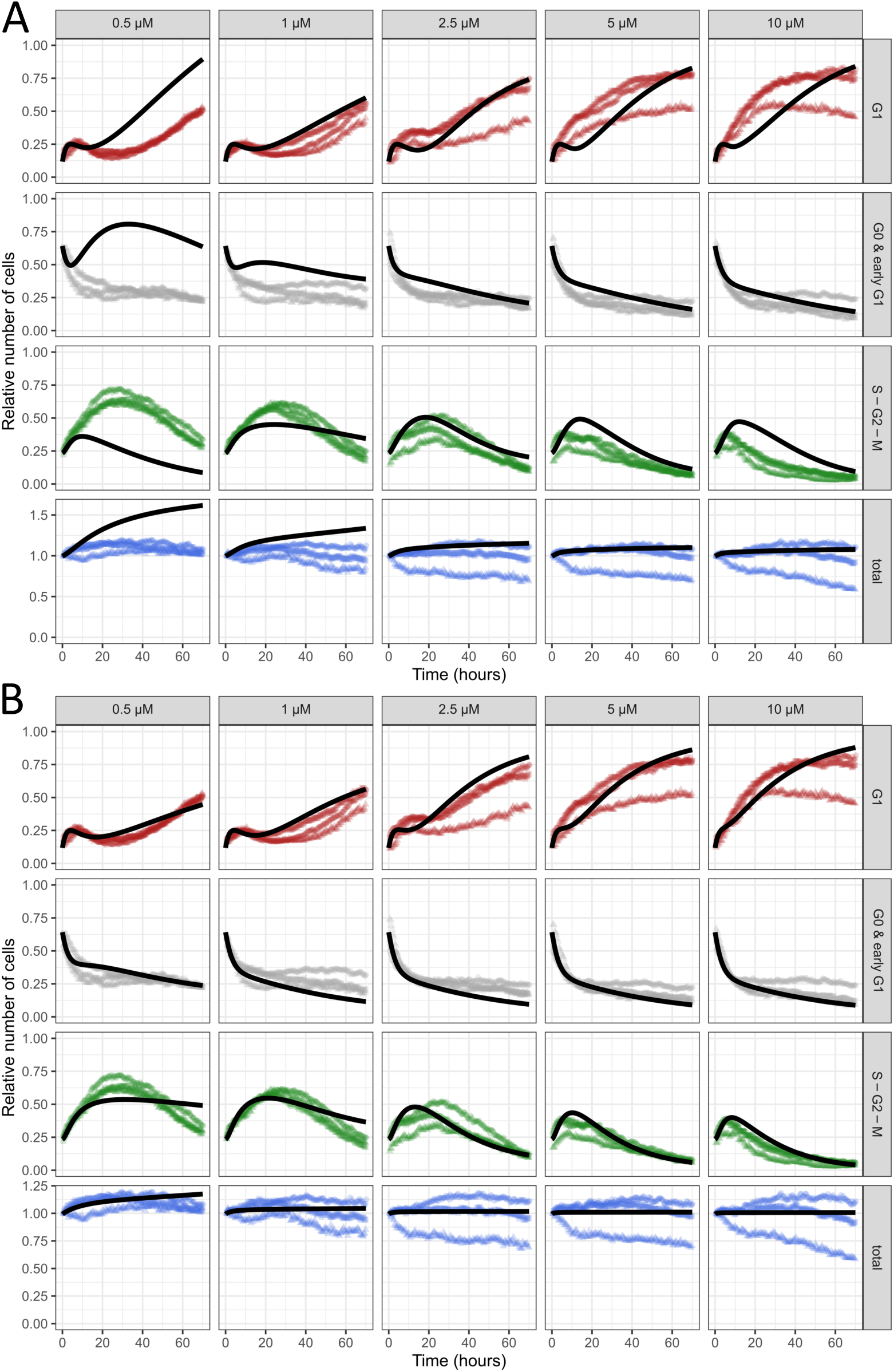
Comparison of cell cycle models for treatment with etoposide. A) Simulations of model without recalibrating any parameters (model M7, solid lines). B) Simulations of model after recalibrating cisplatin-specific parameters (model M9, solid lines). Symbols of different colours represent the three biological replicates.

**Supplementary Figure 8:**
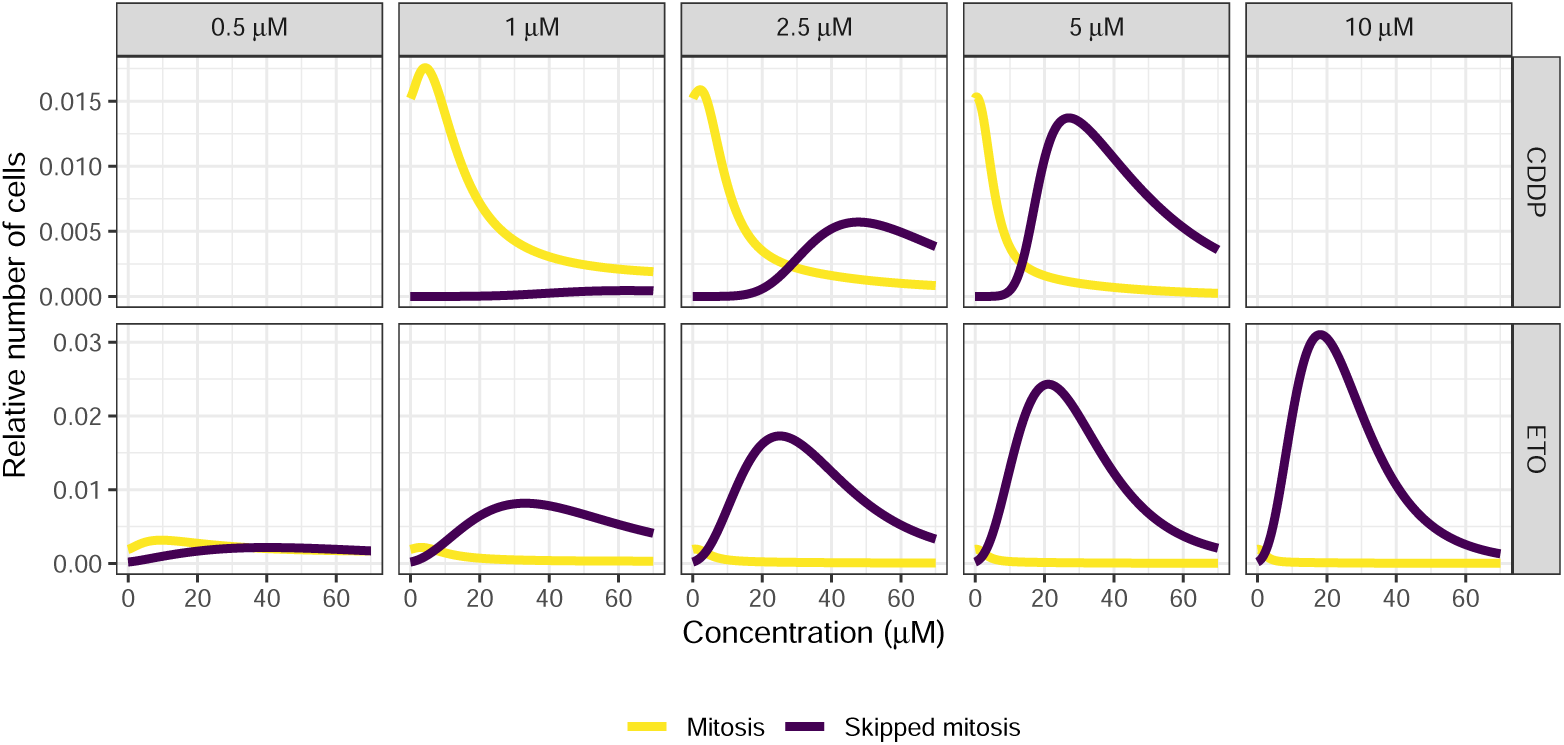
Model prediction of amount of cells undergoing and skipping mitosis over time. Based on model M7 for cisplatin and M10 for etoposide the relative number of cells leaving S-G2-M phase without mitosis (purple) or with mitosis (yellow) was computed for each timestep.

**Supplementary Figure 9:**
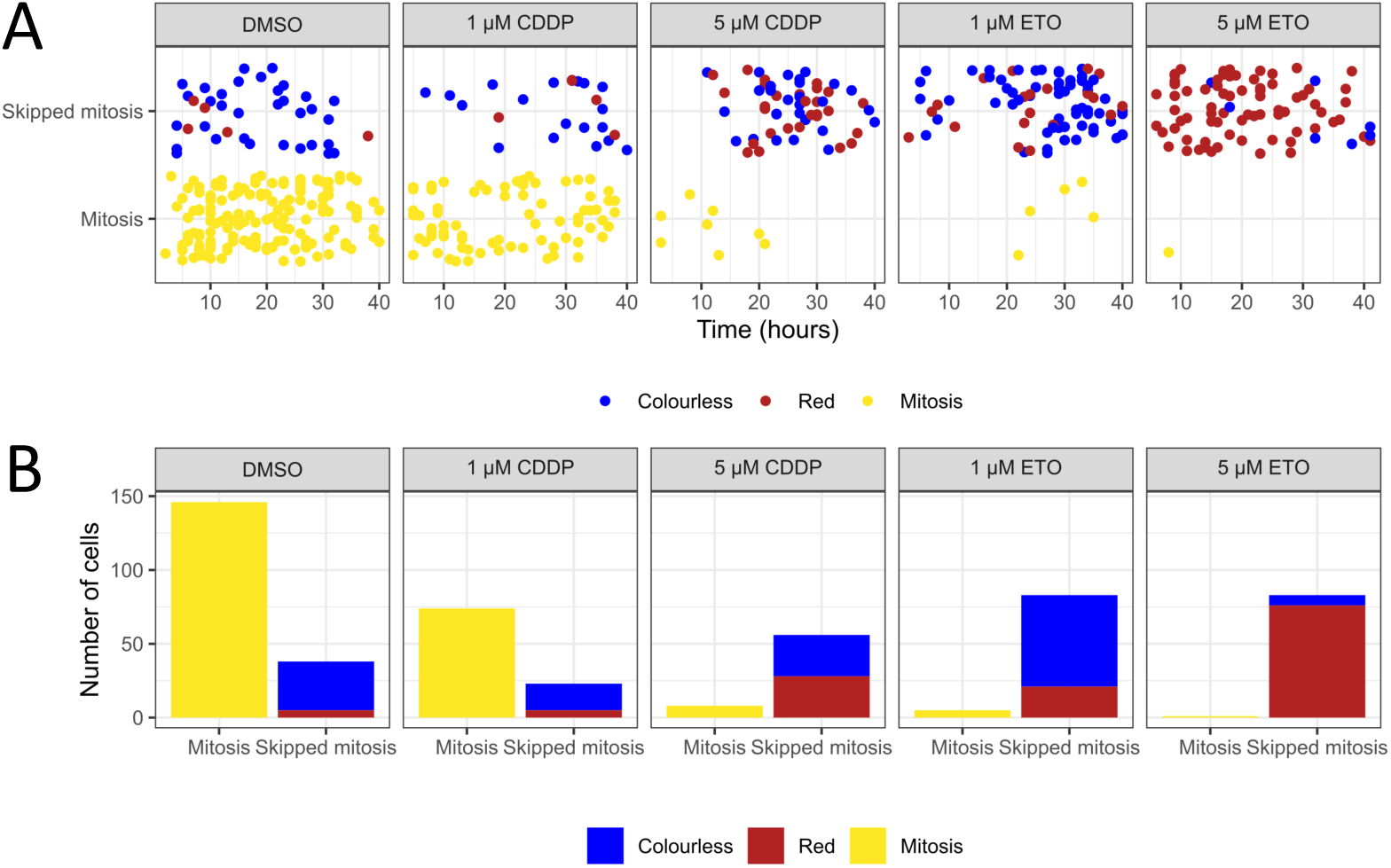
Distribution of colourless and red cells that skipped mitosis in magnified FUCCI data. A) Temporal distribution of cells undergoing and skipping mitosis. Each dot represents the timepoint at which a green cell divided (yellow dots) or transitioned to colourless (blue dots) or red (red dots) without mitosis. B) Number of cells skipping mitosis (red and blue) or undergoing mitosis (yellow). Cells that skipped mitosis are subdivided into cells that were colourless (blue) or red (red) immediately after leaving G2 phase.

### Supplementary Tables

**Supplementary Table 1:** Parameter values for the DDR model. The asterisk (∗) indicates fixed parameter values and the bullet (•) indicates values that were computed with the steady state constraints. For subsequent cell cycle models for etoposide, the values with scaling were used.

| Parameter | Unit | Description | Value CDDP | Value ETO<br>no scaling | Value ETO<br>with scaling |
| --- | --- | --- | --- | --- | --- |
| $EC0$ | - | Initial stress value for 0 $\mu\text{M}$ drug | 0 * | | |
| $EC1$ | - | Initial stress value for 0.5 $\mu\text{M}$ drug | - | 0.1206 | 0.1631 |
| $EC2$ | - | Initial stress value for 1 $\mu\text{M}$ drug | 0.2847 | 0.3503 | 0.4680 |
| $EC3$ | - | Initial stress value for 2.5 $\mu\text{M}$ drug | 0.4791 | 0.7184 | 0.8779 |
| $EC4$ | - | Initial stress value for 5 $\mu\text{M}$ drug | 0.8161 | 1.0314 | 1.2242 |
| $EC5$ | - | Initial stress value for 10 $\mu\text{M}$ drug | - | 1.5754 | 1.6118 |
| $\tau$ | $\text{hr}^{-1}$ | Stress decay rate | 0.01311 | | 0.03339 |
| $\alpha$ | - | Scaling parameter for etoposide | 1 * | | 1.1569 |
| $DD_{init}$ | - | Initial value for DNA damage | 3.4188 | | 3.4188 * |
| $p53_{init}$ | au | Initial value for p53 | 0.09281 | | 0.09281 * |
| $M2_{init}$ | au | Initial value for MDM2 | 0.1371 | | 0.1371 * |
| $p21_{init}$ | au | Initial value for p21 | 0.06098 | | 0.06098 * |
| $B2_{init}$ | au | Initial value for BTG2 | 0.03704 | | 0.03704 * |
| $b_{DD}$ | $\text{hr}^{-1}$ | Basal DNA damage production rate | 0.01231 | | 0.01231 * |
| $b_{p53}$ | au/hr | DNA damage dependent p53 increase rate | 957.81 | | 957.81 * |
| $d_{p53,M2}$ | $\text{au}^{-1}$ | MDM2-mediated p53 degradation rate | 0.000285 | | 0.000285 * |
| $b_{M2}$ | au/hr | Basal MDM2 production rate | 0.01826 | | 0.01826 * |
| $V_{M2}$ | au/hr | Maximal p53-dependent MDM2 production rate | 0.07141 | | 0.07141 * |
| $Km_{M2}$ | au | Michaelis-Menten constant for MDM2 production | 0.2721 | | 0.2721 * |
| $b_{p21}$ | au/hr | Basal p21 production rate | 0.00847 | | 0.00847 * |
| $V_{p21}$ | au/hr | Maximal p53-dependent p21 production rate | 0.04029 | | 0.04029 * |
| $Km_{p21}$ | au | Michaelis-Menten constant for p21 production | 0.4053 | | 0.4053 * |
| $b_{B2}$ | au/hr | Basal BTG2 production rate | 0.08725 | | 0.08725 * |
| $V_{B2}$ | au/hr | Maximal p53-dependent BTG2 production rate | 1.1896 | | 1.1896 * |
| $Km_{B2}$ | au | Michaelis-Menten constant for BTG2 production | 0.4001 | | 0.4001 * |
| $d_{DD}$ | $\text{au}^{-1}\text{hr}^{-1}$ | p53-dependent DNA damage degradation rate | 0.03878 • | | 0.03878 * |
| $d_{p53}$ | $\text{hr}^{-1}$ | Basal p53 degradation rate | 35283 • | | 35283 * |
| $d_{M2}$ | $\text{hr}^{-1}$ | Basal MDM2 degradation rate | 0.1401 • | | 0.1401 * |
| $d_{p21}$ | $\text{hr}^{-1}$ | Basal p21 degradation rate | 0.1407 • | | 0.1407 * |
| $d_{B2}$ | $\text{hr}^{-1}$ | Basal BTG2 degradation rate | 2.448 • | | 2.448 * |

**Supplementary Table 2:** Parameter values for the cell cycle model in control conditions with (model M1) and without (model M2) effect of resources. The asterisk (∗) indicates fixed parameter values. For all subsequent models (M3-M10) the parameters for the model with resources were used.

| Parameter | Unit | Description | Value M1 | Value M2 |
| --- | --- | --- | --- | --- |
| $G1_{early_{init}}$ | - | Initial fraction of cells in early G1 | 0.2924 | 0 |
| $G0_{init}$ | - | Initial fraction of cells in G0 | 0.3495 | 0.3202 |
| $G1_{init}$ | - | Initial fraction of cells in G1 | 0.1175 | 0.1498 |
| $SG2M_{init}$ | - | Initial fraction of cells in SG2M | 0.2337 | 0.5353 |
| $R_{init}$ | - | Initial proportion of resources | 1 * | 1 * |
| $sf$ | - | Scaling factor for depletion of resources | 0.09383 | 0 * |
| $r_{entry}$ | $hr^{-1}$ | Rate of entry from G0 to G1 | 0.3318 | 0.00497 |
| $r_{exit}$ | $hr^{-1}$ | Rate of exit from early G1 to G0 | 0 * | 0 * |
| $r_{G1}$ | $hr^{-1}$ | Rate of transition from early G1 to G1 | 0.01858 | 0.01775 |
| $r_{SG2M}$ | $hr^{-1}$ | Rate of transition from G1 to SG2M | 0.2142 | 0.00103 |
| $r_{mit}$ | $hr^{-1}$ | Rate of mitosis | 0.06511 | 0.05015 |

**Supplementary Table 3:** Parameter values for the cell cycle models without endoreplication. The asterisk (∗) indicates fixed parameter values.

| Parameter | Unit | Description | Value M1 | Value M2 | Value M3 | Value M4 | Value M5 | Value M6 |
| --- | --- | --- | --- | --- | --- | --- | --- | --- |
| $k_1$ | - | Inhibition of mitosis by BTG2 | 67.914 | 119.23 | 67.852 | 68.475 | 0 * | 0 * |
| $k_2$ | - | Inhibition of mitosis by p21 | 0 | 0 | 0 | 0 | 186.67 | 0 * |
| $k_3$ | - | Inhibition of the G1 to S-G2-M transition by BTG2 | 0 | 0 | 1.904 | 0 * | 0 | 180.31 |
| $k_4$ | - | Inhibition of the G1 to S-G2-M transition by p21 | 6.184 | 15.495 | 0 * | 0 * | 3.574 | 296.65 |

**Supplementary Table 4:** Parameter values for the cell cycle models with endoreplication. The asterisk (∗) indicates fixed parameter values.

| Parameter | Unit | Description | Value M7 | Value M8 | Value M9 | Value M10 |
| --- | --- | --- | --- | --- | --- | --- |
| $k_1$ | - | BTG2-mediated inhibition of mitosis rate | 71.209 | 71.209 * | 1000 | 71.209 * |
| $k_2$ | - | p21-mediated inhibition of mitosis rate | 0 | 0 * | 1000 | 0 * |
| $k_3$ | - | BTG2-mediated inhibition of G1 to S-G2-M rate | 0 | 0 * | 7.406 | 0 * |
| $k_4$ | - | p21-mediated inhibition of G1 to S-G2-M rate | 0 | 0 * | 0 | 0 * |
| $r_{end}$ | $hr^{-1}$ | Maximal rate of endoreplication | 0.0390 | 0.0390 * | 0.0514 | 0.3334 |
| $Km_{DD}$ | - | Amount of DNA damage for half-maximal endoreplication rate | 15.169 | 15.169 * | 14.105 | 32.989 |
| $m$ | - | Hill coefficient for endoreplication | 10.00 | 10.00 * | 5.005 | 2.658 |
| $r_{mit}$ | $hr^{-1}$ | Rate of mitosis | 0.06511 * | 0.06511 * | 0.06511 * | 0.00814 |

